# Evidence supporting the first secondary chromosome in actinobacteria as a hallmark of the *Embleya* genus

**DOI:** 10.1101/2025.07.03.662523

**Authors:** Juan Pablo Gomez-Escribano, Siobhan Dorai-Raj, David Baker, Ernest Lacey, Barrie Wilkinson, Thomas J. Booth

**Author notes:** Correspondence. Dr Juan Pablo Gomez-Escribano,; Dr Thomas J. Booth,.

## Abstract

*Embleya* is a genus within the family *Streptomycetaceae*, a group of actinobacteria with outstanding capacity for production of specialised metabolites and a strikingly complex life cycle. In this work, we sequenced the complete genome of the new species *Embleya australiensis* MST-11070 and validated the assembly using optical mapping. The genome of *E. australiensis* MST-11070 consists of a 7.1 Mb linear chromosome and three additional replicons, including a 4.2 Mb linear replicon, EEC1, significantly larger than all previously described secondary replicons from bacteria. EEC1 is typified by its similar composition to the chromosome in terms of GC-content, codon usage and gene functions. It also carries terminal inverted repeats identical to the chromosome. EEC1 is enriched in biosynthetic gene clusters (BGCs), including the only copy of the BGCs for the spore pigment and the surfactant peptide SapB, metabolites essential for the organism’s lifecycle. EEC1 contains an origin of replication with at least some chromosomal properties, and its replication is likely to depend on functions provided by chromosomally located genes. Further comparison of *Embleya* spp. genomes suggests that EEC1-like replicons are conserved across the genus, in contrast to other known large linear extrachromosomal replicons (megaplasmids) in the order. EEC1 is thus a hallmark of the *Embleya* genus and is central to its evolution within the *Streptomycetaceae* family. We propose EEC1 as a secondary chromosome, distinct from previously described secondary chromosomes that utilise plasmid-like replication mechanisms (chromids) and the largest secondary replicon reported in bacteria, to date.

## Introduction

Actinobacteria (Actinomycetota) are a diverse and important phylum of bacteria including major pathogens, rhizobionts and producers of bioactive natural products, including essential medicines. Actinobacteria play host to a wide variety of extrachromosomal elements, namely plasmids^1,2^ and megaplasmids^3–5^. These extrachromosomal replicons play an important role in virulence^6,7^, resistance^8^ and biosynthesis^3^. In addition to harnessing well known producers of bioactive compounds such as *Streptomyces*, exploiting rare and under-represented Actinobacteria is an important strategy for drug discovery^9^. Moreover, exploring these genomes will also lead to breakthroughs in fundamental biology of the phylum.

Within the phylum Actinomycetota is the family *Streptomycetaceae*. Members of this family possess one of the most complex life cycles among bacteria^10,11^. Briefly, monogenomic spores germinate into multigenomic (coenocytic) vegetative mycelium that, in turn, differentiate into aerial hyphae. These hyphae undergo compartmentalisation into the spores as means of reproduction, dispersion and resistance against environment conditions. *Embleya* is a recent genus in the *Streptomycetaceae* family, proposed after the reclassification of *Streptomyces scabrisporus* to *Embleya scabrispora*^12,13^, the type-species of the genus, in 2018^12^. At the time of writing, the genus officially contains only one other species, *Embleya hyalina* (as stated in the LPSN database^14^, last accessed on 8^th^ October 2024). *E. scabrispora* and *E. hyalina* are known to produce the specialised metabolites hitachimycin^15,16^ and nybomycin^17,18^, respectively. Several additional strains have been identified by 16S sequencing, but taxonomic descriptions are poor, with *Embleya* strains being erroneously described as *E. scabrispora*^19^. The NCBI’s refseq database contains few *Embleya* genomes, most of which are relatively low-quality assemblies (N50 < 1 Mb). However, during this study, two additional high-quality sequences were published alongside over 1,000 additional actinobacterial genomes^20^, but the characterisation of these strains was limited. A recent bioinformatic analysis has revealed the diversity and breadth of biosynthetic gene clusters (BGCs) present in *Embleya* genomes^19^. However, the lack of complete *Embleya* genomes severely restricts the possible analyses. Given the potential value of *Embleya* spp. there is significant need for high quality *Embleya* genomes to provide a foundation for exploring this enigmatic genus.

Here, we describe the genome of a new species, *Embleya australiensis* MST-111070, a known producer of nybomycins, leptomycins, kazusamycins and Antibiotic L-1566027. We describe the presence of a giant 4.2 Mb secondary replicon and confirm its existence through optical mapping. Finally, we conduct an in depth bioinformatic analysis of the secondary replicon and provide evidence for an essential role during the evolution of the genus, leading to the proposal for the classification of this replicon as a secondary chromosome.

## Results and Discussion

### *Embleya australiensis* MST-111070 contains a 4.2 Mb second replicon

#### The complete genome of *Embleya australiensis* MST-111070

Strain MST-111070 was isolated by the biotechnology company Microbial Screening Technologies (Smithfield, Australia) as part of a programme to identify talented, free-growing Actinomycetes from arid soils. Agar cultivation of the strain appeared morphologically as a typical streptomycete that on methanolic extraction and LCMS showed high level production of several rare, specialised metabolites: nybomycin and deoxynybomycin; leptomycin A and B and kazusamycin A and B; and antibiotic L 156602 (Microbial Screening Technologies’ BioAustralis metabolite catalogue and unpublished data). Following the genomic sequencing performed as part of this work, and the taxonomic analysis detailed below, we have reclassified the strain as *Embleya australiensis sp. nov.*.

High molecular weight DNA was extracted from *E. australiensis* MST-111070 and sequenced with Pacific Biosciences RSII SMRT technology (PacBio) at the Earlham Institute (Norwich, UK). In total, we attained over 900 Mb of raw sequence data, corresponding to approximately 90x coverage of an average *Streptomyces* genome. The genome was assembled using HGAP2 and HGAP3 independently (Table 1). Both assemblies were essentially equivalent, consisting of three contigs of 7.1, 4.2 and 0.2 Mb. The HGAP3 assembly included an extra contig of 0.3 Mb. It must be noted that both assemblies originate from the same raw data, therefore any difference is accounted for exclusively by the changes in the algorithms from version 2 to 3 of the assembly pipeline.

**Table 1:** Comparison of the assemblies generated in this study. The size of the *Embleya australiensis* MST-111070 chromosome and each *<u>E</u>mbleya* <u>E</u>xtrachromosomal <u>E</u>lement (EEC1-3) is shown in bp. Version 4 was submitted to NCBI.

| Assembly version | Method | Chromosome | EEC1 | EEC2 | EEC3 | Total |
| --- | --- | --- | --- | --- | --- | --- |
| 1 | Pacific Biosciences RSII, HGAP2 | 7,100,744 | 4,187,648 | - | 228,860 | 11,517,252 |
| 2 | Pacific Biosciences RSII, HGAP3 | 7,101,142 | 4,187,746 | 302,592 | 228,392 | 11,820,375 |
| 3 | Illumina MiSeq, University of Cambridge | 7,075,274<br>21,156 | 4,164,320 | - | 207,392 | 11,468,142 |
| 4 | Pacific Biosciences RSII and Illumina consensus | 7,117,762 | 4,217,944 | 302,592 | 210,636 (circular) | 11,848,934 |

The presence of the large 4.2 Mb contig led us to consider that our assembly might not accurately represent the genome. Therefore, we sought to obtain an independent *de novo* assembly using a different sequencing technology. To achieve this, we turned to Illumina MiSeq using paired-end sequencing which was, crucially, combined with a Nextera Mate Pair shotgun library and the in-house assembly pipeline at the University of Cambridge DNA sequencing facility (Department of Biochemistry)^21^. This approach produced an assembly consisting of a set of contigs that were very similar to that obtained with PacBio HGAP2, including the 4.2 Mb contig (Table 1).

Subsequently, the contigs from both PacBio assemblies were aligned with the Illumina contigs, and the alignment was manually curated. The Illumina contigs also allowed the extension of the ends of the PacBio contigs (for similar strategy see^22,23^) to generate a final combined assembly of four contigs: 7.1 Mb and 4.2 Mb of linear topology; 0.3 Mb apparently linear according to optical mapping, see below; and 0.2 Mb of circular topology (Table 1). These contigs representing the chromosome and the extra-chromosomal elements (here designated ***<u>E</u>****mbleya* **<u>e</u>**xtra**<u>c</u>**hromosomal elements, EEC1 to 3) were submitted to GenBank with accessions CP182422 - CP182425.

#### Terminal inverted repeats in the *Embleya australiensis* MST-111070 genome

In addition to linear chromosomes, many other members of the *Streptomycetaceae* family host additional linear extrachromosomal replicons, all of which contain characteristic terminal inverted repeats (TIRs)^22,24^. TIRs are large regions of DNA (typically 20 Kb – 1 Mb)^1,20,25^ found at both ends of the replicon and are required for the replication and stability of linear replicons^26–29^. The presence of TIRs in an assembly is usually indicative of a nearly complete replicon sequence. Analysis of the 7.1 Mb contig revealed the presence of terminal inverted repeats of around 21.2 kb. Interestingly, the TIRS of the 4.2 Mb replicon are almost identical to those of the chromosome (99.95 % identity). Furthermore, the homology between the right ends of the genome and EEC1 extends to 33.6 kb of sequence with 99.94% identity (see Figure 1 and Figure S1 for further details). This extraordinary share of replicon-end asymmetry could be explained by a hypothetical origin of EEC1 as a schism event from the chromosome, followed by homogenisation among the ends of both replicons, a commonly observed mechanism for the maintenance of TIRs^27,29^. The presence of TIRs in EEC1 is strong evidence that it is a standalone replicon, but, given the extraordinary size of EEC1, we sought further independent evidence to confirm its existence and structure as assembled by both DNA sequencing technologies.

**Figure 1:**
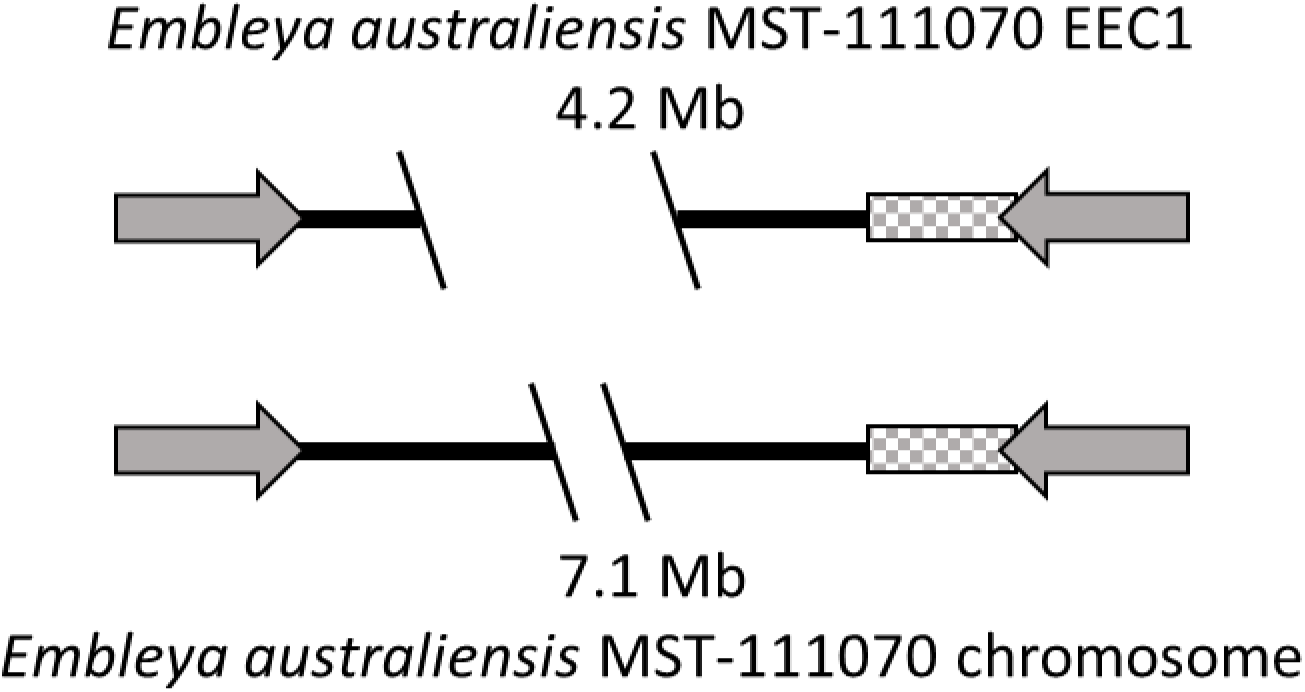
Relationship between the *Embleya australiensis* MST-111070 chromosome and EEC1. The 21.2 kb terminal inverted repeats (TIRs) are shown as solid grey arrows, and the 12.4 kb block of additional shared homology is shown as grey checked bar.

#### Confirmation of genome assembly and replicon topology with optical mapping

The genome assembly suggested the existence of a large (4.2 Mb) secondary replicon which, if confirmed, would be the largest secondary replicon reported from actinobacteria and of a size comparable to the *E. coli* chromosome. With whole genome sequencing, there is always the possibility that the sequence is misassembled into artefactual contigs that are not representative of the actual genome structure^30^, which could be responsible for the apparent split of the expected chromosome into two contigs. Given the unique nature of the EEC1 replicon, we sought to confirm these results through an independent method.

Optical mapping is a technology that produces high resolution genome maps of specific sequence motifs recognised by nickase enzymes. These sites are fluorescently labelled and can be mapped by subsequent imaging of individual DNA molecules in a linear ordered fashion. Therefore, optical mapping can provide an accurate map of nickase sites distribution along individual DNA molecules. These molecules can be used for a *de novo* assembly into contigs representing a consensus linear distribution of nickase sites. Alternatively, the contigs or the individual molecules can be aligned to the predicted distribution of nickase sites along a reference DNA sequence. Because of the large size of the molecules (with sizes exceeding 500 kb in some cases) optical mapping provides a highly accurate and reliable way of checking the accuracy of a DNA sequence assembly and to resolve difficult repeating regions like the TIRs^22,31,32^. We previously showed the application of this technique to confirm an assembly of the *Streptomyces clavuligerus* ATCC 27064 genome^22^.

Here we employed Bionano optical mapping technology to obtain a high-resolution map of the recognition site for nickase Nt.BspQI (GCTCTTC) along the genome. The analysis accounted for 229509 molecules totalling 37288.4 Mb with a molecule N50 of 158.5 Kb, and over 2000 molecules larger than 500 kb. We used the data to confirm the DNA assembly through two independent methods: i) an alignment of Bionano molecules to the Illumina-PacBio hybrid assembly; and ii) generation of a reference-independent *de novo* assembly of Bionano molecules. Firstly, the molecules-to-reference alignment generated statistics indicators that supported an overall reliable alignment of Bionano molecules to the DNA assembly (Map Rate, 42.7%, FP 0.48/100kbp, FP 4.4%, and FN 18.9%; see Materials and Methods and Supporting Information for details and documentation on interpretation of results). The molecules aligned with very high confidence, including along the TIRs (see Figures S2 to S4) providing strong support of the 4.2 Mb contig being a true replicon independent from the 7.1 Mb chromosome. Second, the Bionano *de novo* assembly was performed without the reference genome and the resulting assembly consisted of 8 contigs spanning a total of 14359 kb, with N50 of 4330 kb and an average coverage of 106.3 molecules. 5 contigs mapped to the chromosome while the remaining 3 mapped to each of the extrachromosomal elements; in all cases the DNA contigs were fully covered by the Bionano contigs (see Extended Methods in Supporting Information and Figures S3 and S4). The ends of the DNA contigs were fully supported by the Bionano *de novo* assembled contigs. In addition, the contig matching EEC3 supported the proposed sequence and circular topology (Figure S4). Furthermore, the Bionano analysis confirmed the existence of EEC2, which was previously not assembled in the Illumina and the HGAP2 assemblies and supports a linear topology of this replicon (Table 1, Figure S3). Taken together, the DNA sequencing data and optical mapping confirm the genome of *E. australiensis* MST-111070 as follows: a 7.12 Mb linear chromosome; two plasmids: one large circular plasmid (211 kb, EEC3), another large linear plasmid (303 kb, EEC2); and an extraordinary 4.2 Mb linear replicon (EEC1).

### Taxonomy of *Embleya australiensis* MST-111070

Phylogenetic analysis of the genome of MST-111070 indicated this strain is potentially a new species within the *Embleya* genus, for which we propose the species epithet *E. australiensis*. Firstly, analysis of the genome of *E. australiensis* MST-111070 with the Type Strain Genome Server^33^ (TYGS; first time on 2020-12-28) revealed that its closest relatives are *Embleya hyalina* NBRC 13850T and *Embleya scabrispora* DSM 41855 (digital DNA hybridisation (dDDH d4) scores of 54.8% and 30.9% respectively). TYGS analysis also highlighted the close relationship between the *Embleya* and *Yinghuangia* genera (Figure S5). As the TYGS database contains only genomes of type strains, and it is not constantly updated, we performed an extended analysis adding in any genomes identified in NCBI databases as being possibly related to strain MST-111070 using both TYGS and GGDC^33,34^. This gave additional reassurance that this strain is a member of the genus *Embleya*, which is closely related to *Yinghuangia* (Figure S6). However, there were some discrepancies between the methods, most notably that in the extended 16S analysis, *Yinghuangia* was paraphyletic (Figure 6B). Additionally, *Parastreptomyces abscessus* clades within *Yinghuangia* and *Streptomyces lasii* clades within *Embleya*. To preserve the monophyly of *Embleya*, *Streptomyces* and *Yinghuangia*, these strains should be reassigned to *Yinghuangia* and *Embleya*, respectively.

To clarify the phylogenetic relationship within the *Streptomycetaceae* family and other actinobacteria genera, we undertook an independent and extensive analysis with getphylo^35^ to build a genome-scale phylogenetic tree from 515 complete actinomycete genomes (Figure 2). This phylogeny supports the placement of *E. australiensis* MST-111070 within the genus *Embleya* and supports *Embleya* as a monophyletic group within the *Streptomycetaceae* family, among strains with high quality sequenced genomes. It also confirms *Yinghuangia* as a monophyletic sister genus.

**Figure 2:**
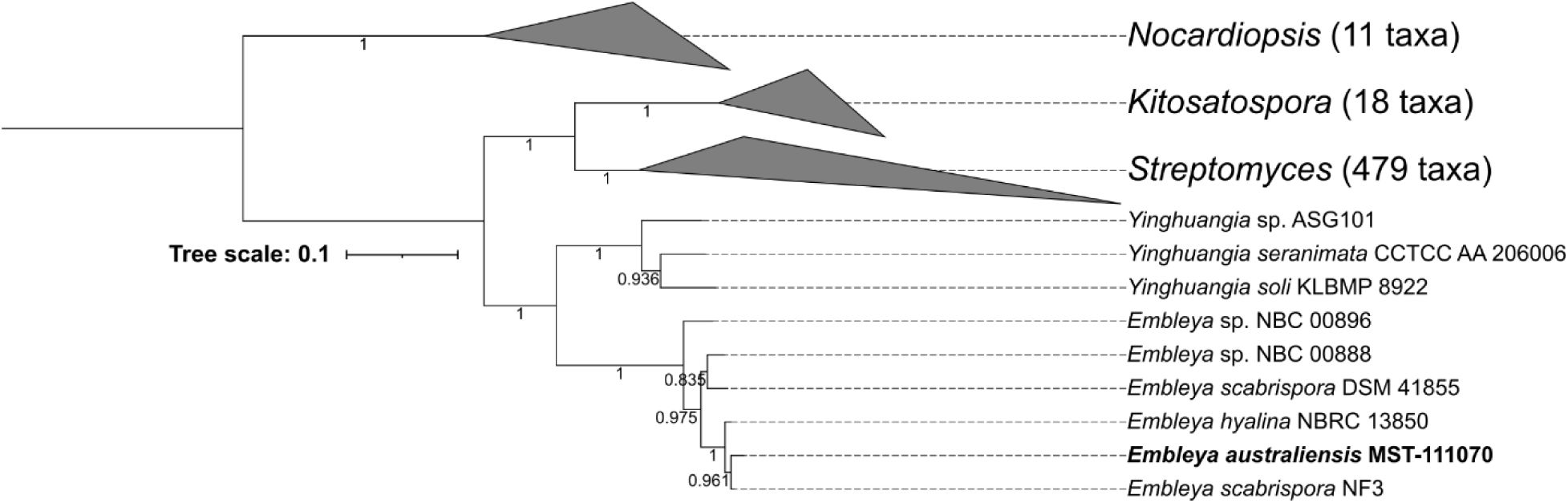
Phylogenetic tree of *Streptomycetaceae*. A phylogenetic tree of 515 strains inferred from 11 loci (Table S1) by getphylo^35^ using Fasttree2^78^ and visualised in iTOL^77^. Clades for *Nocardiopsis* (outgroup), *Kitosatospora* and *Streptomyces* have been collapsed to highlight the inter-generic relationships, especially between *Yinghuangia* and *Embleya*. The tree shows that each genus is monophyletic with maximum support. Branch supports are approximate maximum likelihoods.

### EEC1: Secondary chromosome, chromid, or megaplasmid?

There has been much discussion over the distinct types of bacterial replicons^2,36^. Bacterial genomes are typified by a singular linear or circular chromosome. Many bacteria have additional replicons that are typically termed plasmids or megaplasmids. The distinction between plasmids and megaplasmids is solely based on size. Generally, a plasmid is considered a megaplasmid if it is >350 kb in size^2,36^. Plasmids and megaplasmids are ‘non-essential’ replicons that differ from the chromosome in function and composition. A third type of replicon, the secondary chromosome has been described in specific bacterial lineages, most notably: *Agrobacterium*^37^, *Burkholderia*^38,39^, *Pseudoalteromonas*^40,41^, *Rhodobacter*^42^ and *Vibrio*^43^. An alternative name, ‘chromid’, was proposed by Harrison *et al*., in 2010^36^. Chromids share features of both plasmids and chromosomes; ‘not a chromosome, not a plasmid.’ Most importantly, they replicate using a plasmid-like mechanism. It is hypothesised that chromids have evolved from plasmids but, over long evolutionary time frames, have become essential through the transfer of core genes from the chromosome^36^. Chromids, therefore, demonstrate conservation on a higher taxonomic order and are typically conserved across the genus. An additional distinction has been proposed in which ‘secondary chromosome’ only denotes a replicon that has arisen though a schism of the chromosome^2^. However, for reasons outlined below, this distinction is problematic and, in some fields, the term secondary chromosome is still rarely used or used synonymously with chromid^38,44,45^. For clarity, the features of plasmids, chromids and chromosomes are listed in Table 2.

**Table 2:** Distinguishing features of chromosomes, secondary chromosomes, chromids, and megaplasmids. Adapted from Harrison *et al*.,^36^ and DiCenzo *et al*.^2^ and does not include the observations in this study. Relative terms are given as objective measures vary wildly between bacterial phyla.

| Feature | Megaplasmid | Chromid<br>(Secondary Chromosome) | Secondary Chromosome | Chromosome |
| --- | --- | --- | --- | --- |
| Observed in bacteria | Yes | Yes | No | Yes |
| Core Genes | No | Yes | Yes | Yes |
| Nucleotide Composition | Variable | As chromosome | As chromosome | Variable |
| Codon Usage Bias | Average | High | Highest | Highest |
| Replication System | Plasmid | Plasmid | Chromosomal | Chromosomal |
| Gene Origins | Strain-specific | Genus-specific | Genus-specific | Less genus specific |
| Evolutionary Origin | Plasmid | Plasmid | Chromosome | Chromosome |

#### Replicon size

The first and most apparent feature of any replicon is its size. The chromosome is universally the largest replicon in the bacterial genome. Chromids, when present, are the second largest, i.e., larger than any plasmid. EEC1 is fourteen times larger than the next biggest plasmid, EEC2. EEC1 is nearly 60 % the size of the chromosome and accounts for 35.5% of the total genome. This clear distinction in size makes EEC1 larger than the typically agreed size for megaplasmids^2,36^ (<2 Mb) and into the realm of chromids and chromosomes.

#### Nucleotide composition and codon usage

Chromids and secondary chromosomes share similar GC-content and codon usage with the chromosome, whereas plasmids typically have different GC content (lower in the case of *Streptomycetaceae*)^2,35^. We clearly observe this pattern in the genome of *E. australiensis* MST-111070. The chromosome and EEC1 share almost identical GC contents of ∼71.6 % compared to 69.1 % and 69.5 % for EEC2 and EEC3, respectively. The codon usage in EEC1 is also very close to that of the chromosome. In addition, the chromosome and EEC1 show a greater degree of codon bias than the plasmids, which are more likely to utilise rare codons (Figure 3; Figure S7).

**Figure 3:**
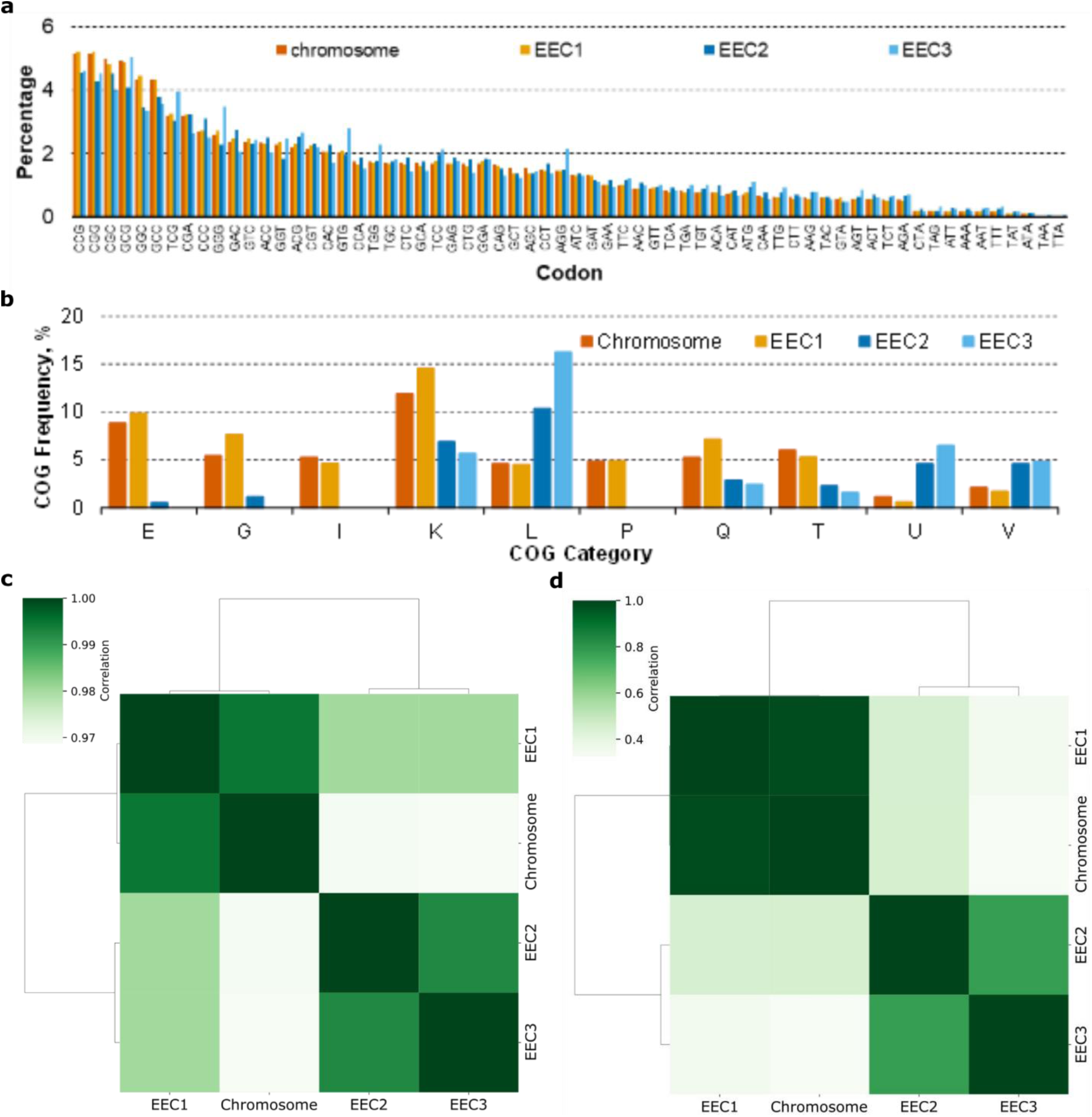
Biochemical and functional composition of *Embleya australiensis* MST-111070 replicons. Bar charts show the percentage abundance of (**a**) selected codons and (**b**) selected COG categories (E: amino acid metabolism and transport; G: carbohydrate metabolism and transport; I: lipid metabolism; K: transcription; L: replication and repair; P: inorganic ion transport and metabolism; Q: secondary metabolites biosynthesis, transport, and catabolism; T: signal transduction; U: intracellular trafficking and secretion; and V: defence mechanisms) across the chromosome and the three extrachromosomal elements (EEC1 - EEC3). Heatmaps show the spearman’s rank correlation for (**c**) the codon usage (calculated with codoniser, this study) and (**d**) the COG category abundance (calculated with egger, this study).

#### Replicon function and the presence of core genes

A defining feature of plasmids is that they are nonessential. Meanwhile, chromids and secondary chromosomes contain core genes and are believed to be essential. Analysis of the genome with the BUSCO (Benchmarking Universal Single-Copy Orthologs)^46^ pipeline revealed that EEC1 contained a significant number of BUSCOs duplicated with the chromosome (Table 3). However, EEC1 also carried 27 unique orthologues that are universally present in *Streptomyces* spp. suggesting that it may be essential.

**Table 3.** Comparative analysis for assembly completeness using BUSCO. The number of BUSCOS indicated for each database correspond to running the pipeline with the full assembly (bold) versus the assembly lacking the 4.2 Mb EEC1. Results indicate that EEC1 carries several duplicated BUSCOs, but also that without this replicon the genome is missing 27 genes typically present in *Streptomycetales* species.

|  | streptomycetales | actinobacteria_<br>class | actinobacteria_<br>phylum | bacteria |
| --- | --- | --- | --- | --- |
| Complete BUSCOs | **1453**/1427 | **356**/356 | **292**/292 | **124**/124 |
| Complete and single-copy BUSCOs | **1427**/1412 | **348**/351 | **286**/288 | **120**/122 |
| Complete and duplicated BUSCOs | **26**/15 | **8**/5 | **6**/4 | **4**/2 |
| Fragmented BUSCOs | **11**/10 | **0**/0 | **0**/0 | **0**/0 |
| Missing BUSCOs | **115**/142 | **0**/0 | **0**/0 | **0**/0 |
| Total BUSCO groups searched | **1579**/1579 | **356**/356 | **292**/292 | **124**/124 |

We also compared functional content of the replicons by comparing the frequency of functionally annotated COGs (Clusters of Orthologous Genes). We annotated the *E. australiensis* MST-111070 genome using eggNOG^47^ as implemented in eggNOG-mapper^48^. The analysis revealed that EEC1 is functionally more similar to the chromosome than the plasmids, whereas the plasmids are more alike (Figure 3; Figure S8). The chromosome and EEC1 are enriched in metabolic functions (E, G, I, P, Q), and functions associated with transcription (K) and signal transduction (T). On the other hand, the two smaller replicons are enriched in genes for replication and repair (L), intracellular trafficking, secretion and vesicular transport (U), and defence (V). They also contain a much higher proportion of genes with unknown or unassigned functions. This disproportion may be due to size, since smaller replicons will dedicate a proportionally higher percentage of genes to maintenance and replication). However, the complete lack of some COGs from the plasmids, namely E (amino acid transport and metabolism), I (lipid transport and metabolism) and P (inorganic ion transport and metabolism) shows a definitive functional bias. These COGs are each associated with primary metabolic functions and suggests that the EEC1 has a much more central role in metabolism than either of the plasmids.

Finally, EEC1 is enriched in biosynthetic gene clusters (BGCs). BGCs are co-located groups of genes responsible for the biosynthesis of small molecules with specialised functions such as antibiotics and siderophores^49,50^. Analysis of the *E. australiensis* MST-111070 genome with antiSMASH 7.0^50^ predicted a total of 49 biosynthetic regions (Table S3; Figure S9). Surprisingly, the majority of these regions (25) are found on EEC1 and not the chromosome (24). The plasmids contain a single BGC each. For its size EEC1 shows a remarkable density of specialised biosynthetic machinery, almost double that of the chromosome. Although the majority of the BGCs encode the biosynthesis of unknown products (cryptic pathways), there are several BGCs that can be assigned to molecules known to be produced by this strain (Table 4). Antibiotic L 1566025 was assigned to a hybrid polyketide-non ribosomal peptide synthase BGC due to the domain organisation of the modular megasynthases and its similarity to the BGC of the related acylated cyclic hexadepsipeptide polyoxypeptin^51^. Nybomycin was assigned to a small BGC with high similarity to the nybomycin BGC previously described^18^. Finally, leptomycins and kazusamycin were assigned to a polyketide BGC that showed high nucleotide identity (92 %) and identical organisation to the BGC described in patent US7288396B2^52^. The biosynthesis of the known products of *E. australiensis* MST-11080 is restricted to the chromosome (nybomycin and leptomycins and L-156602) although the biological significance of this observation, if any, is unclear.

**Table 4.**
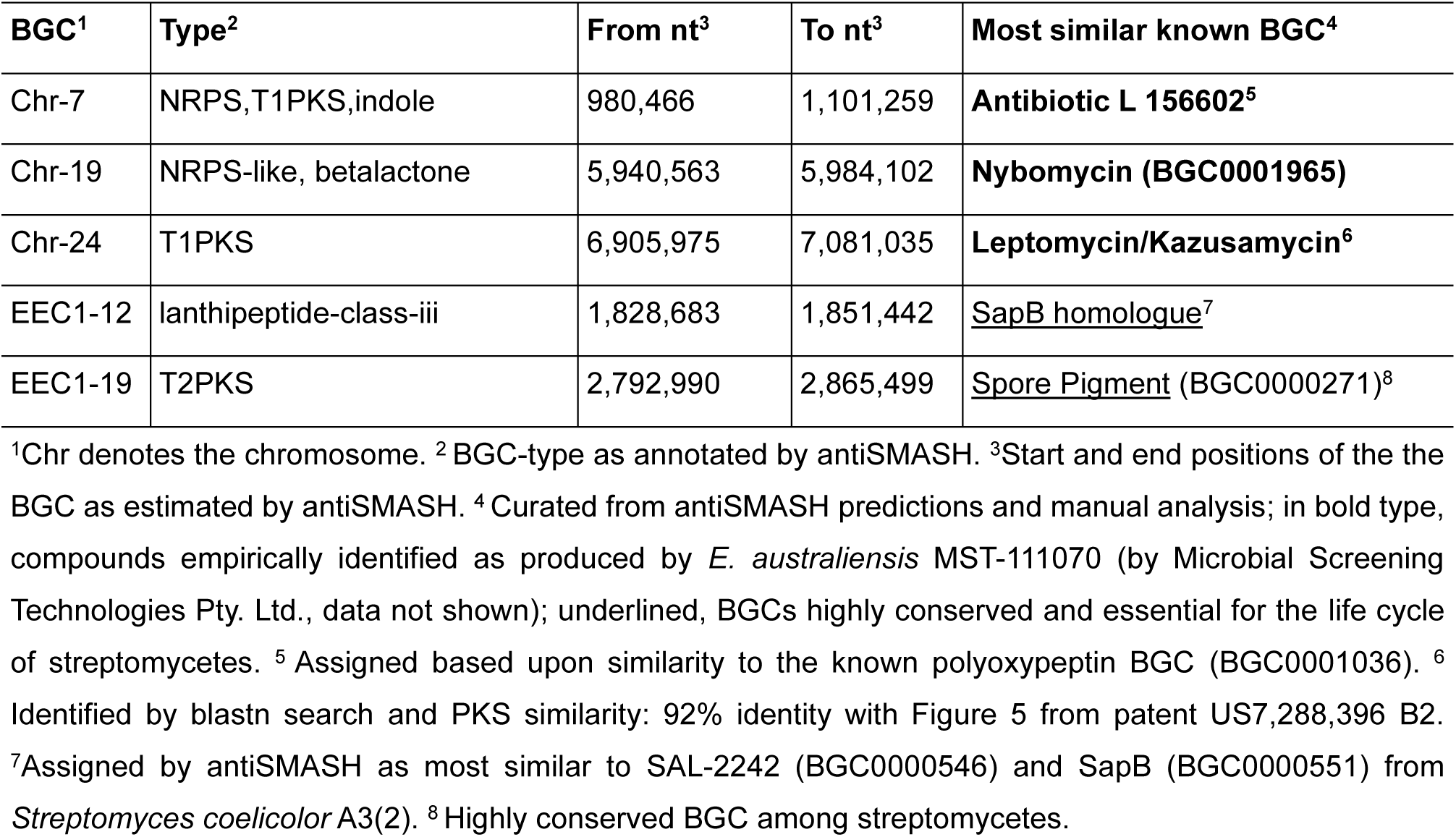
Selected BGCs from a curated antiSMASH output for the genome of *Embleya australiensis* MST-111070.

| BGC<sup>1</sup> | Type<sup>2</sup> | From nt<sup>3</sup> | To nt<sup>3</sup> | Most similar known BGC<sup>4</sup> |
| --- | --- | --- | --- | --- |
| Chr-7 | NRPS,T1PKS,indole | 980,466 | 1,101,259 | **Antibiotic L 156602<sup>5</sup>** |
| Chr-19 | NRPS-like, betalactone | 5,940,563 | 5,984,102 | **Nybomycin (BGC0001965)** |
| Chr-24 | T1PKS | 6,905,975 | 7,081,035 | **Leptomycin/Kazusamycin<sup>6</sup>** |
| EEC1-12 | lanthipeptide-class-iii | 1,828,683 | 1,851,442 | <u>SapB homologue<sup>7</sup></u> |
| EEC1-19 | T2PKS | 2,792,990 | 2,865,499 | <u>Spore Pigment (BGC0000271)<sup>8</sup></u> |

<sup>1</sup>Chr denotes the chromosome. <sup>2</sup> BGC-type as annotated by antiSMASH. <sup>3</sup>Start and end positions of the the BGC as estimated by antiSMASH. <sup>4</sup> Curated from antiSMASH predictions and manual analysis; in bold type, compounds empirically identified as produced by *E. australiensis* MST-111070 (by Microbial Screening Technologies Pty. Ltd., data not shown); underlined, BGCs highly conserved and essential for the life cycle of streptomycetes. <sup>5</sup> Assigned based upon similarity to the known polyoxypeptin BGC (BGC0001036). <sup>6</sup> Identified by blastn search and PKS similarity: 92% identity with Figure 5 from patent US7,288,396 B2. <sup>7</sup>Assigned by antiSMASH as most similar to SAL-2242 (BGC0000546) and SapB (BGC0000551) from *Streptomyces coelicolor* A3(2). <sup>8</sup> Highly conserved BGC among streptomycetes.

Strikingly, among the BGCs encoded by EEC1 are both the BGC required for biosynthesis of the spore pigment and a SapB homologue. Both BGCs play an important role in the morphological development and life cycle of *Streptomyces* spp. and related Actinobacteria: spore pigment is found in mature spores, whereas SapB-like peptides play a vital role in the development of aerial mycelium^53,54^. Again, the presence of these BGCs on EEC1 supports the conclusion that it encodes essential cellular functions.

#### Replicative mechanism

Despite their overall similarity to the chromosome, chromids utilise plasmid-type replication systems^2,36^. In *Streptomyces* species, linear plasmids commonly replicate bidirectionally from a central replicon consisting of a DNA helicase (*rep2*) and it’s corresponding iteron (*rep1*)^55–57^. In contrast, the chromosomal origin of replication is flanked by the replication initiation protein, *dnaA* and the DNA polymerase subunit, *dnaN*^58–60^. Both replication systems require a set of partitioning genes *parA* and *parB*^5,61,62^. The origin of each *E. australiensis* MST-111070 replicon were identified using Diamond homology searches against previously described replicons^5,62^ (Table S2 and Figure 4). The chromosomal origin was easily identified by the presence of *dnaA* and *dnaN* homologues in addition to homologues of *parA*, *parB*, *gyrA, gyrB* in the local regions (loci: ACMXN5_16320 - ACMXN5_16385). The origin of EEC3 was also easily identified with homology searching due to the presence of close homologues of *rep1* and *rep2* (ACMXN5_50245 and ACMXN5_50250) and *parA* and *parB* (ACMXN5_50205 and ACMXN5_50210). The replication systems of EEC1 and EEC2 are much more difficult to define. EEC1 lacks the canonical plasmid replication system, but it also lacks genes required for chromosomal replication. It carries a copy of *dnaN* (ACMXN5_39850) but is lacking a corresponding *dnaA* locus. A distant homologue of *parA* is also present in the vicinity (ACMXN5_39820). Further searching using PFAM HMMs also failed to find *parB* and *dnaA* homologues. One possibility is that the replication of EEC1 relies on machinery encoded in the chromosome, i.e. it cannot replicate independently from the chromosome. EEC1 may also replicate through an, as yet, unidentified mechanism. EEC2 has both *parA* and *parB* homologues (ACMXN5_49550 and ACMXN5_48975), however they are not clustered. Additionally, although EEC2 encodes several helicases, none show homology with helicases from canonical plasmid or chromosomal replicons. Overall, these results highlight the diversity of replicons in *Streptomycetaceae* that have yet to be characterised. GC-skew analysis was consistent with the location of the chromosomal origin and supported that EEC3 is a circular plasmid. The GC-skew of EEC1 shows a plateau between 1.9 and 2.8 Mb, which may indicate a recent rearrangement.

**Figure 4.**
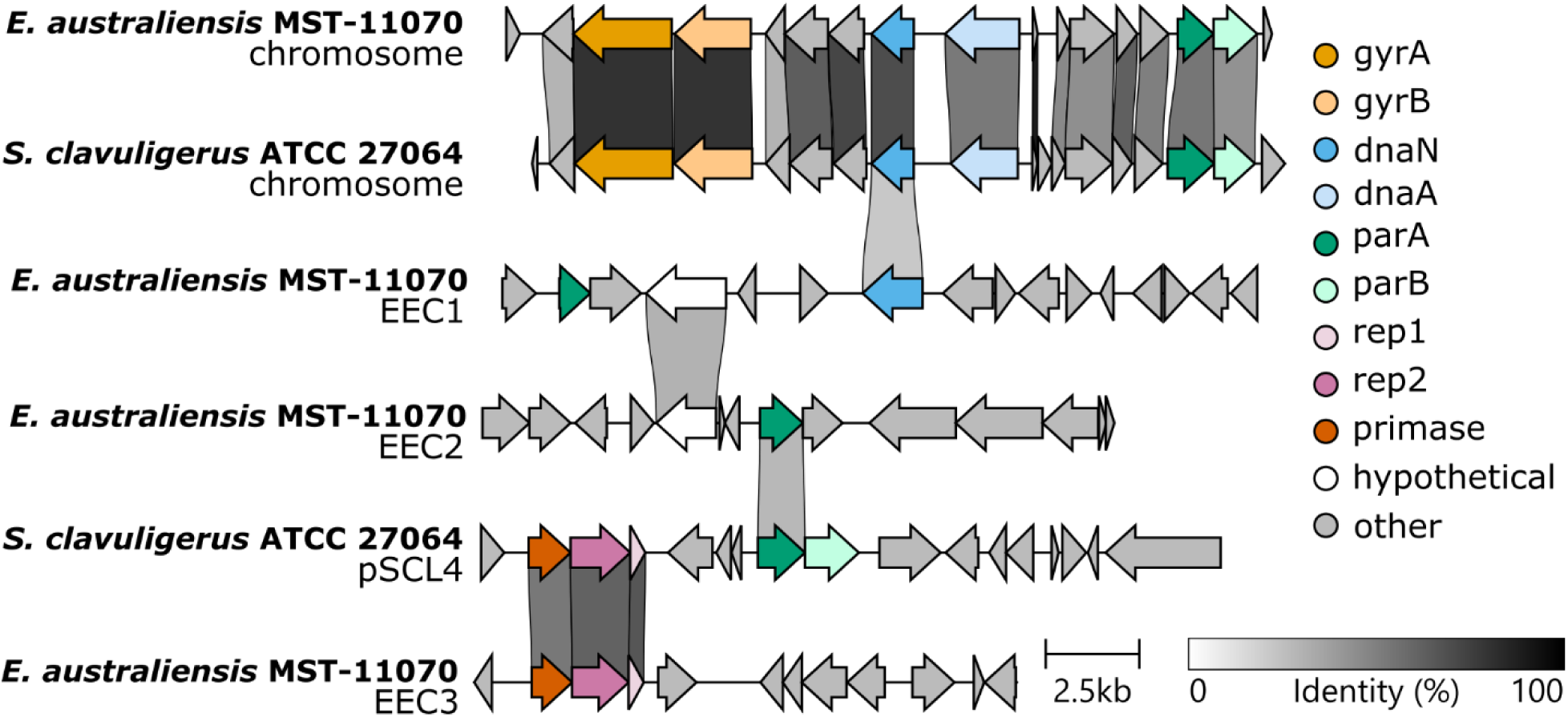
Comparison of predicted replicon regions carrying the origin of replication. Modified from clinker output^79^. Links show genes encoding proteins sharing an identity of >30 %. Genes are coloured by the predicted function based on profile HMMs^80^.

#### Evolutionary origin and genus specificity

Another hallmark feature of chromids is that they contain a high number of genus-specific genes (i.e. genes that only have close homologues within the genus)^36^. We thus estimated the specificity of genes on the strain-, genus- and family-level (Figure S10-S12). In general, the chromosome shows a trend of increasing specificity with higher taxonomic rank (i.e. lower proportion of strain-specific genes), whereas the plasmids EEC2 and EEC3 show the inverse (higher proportion of strain-specific genes). At all taxonomic ranks, EEC1 shows similar specificity to the chromosome, exhibiting highest levels of specificity at the strain and genus level. While most of the genes on EEC2 and EEC3 are unique to those plasmids, EEC1 encodes genes that are also encoded by other *Embleya* strains. This indicates a high level of conservation of EEC1 across *Embleya* species, in line with the chromosome.

While preparing this manuscript, high quality genome assemblies for two additional *Embleya species* were published from the New Bioactive Compounds (NBC) collection^20^. These genome assemblies also contain large secondary replicons (3.1 Mb and 5.7 Mb) that exhibit similar traits to EEC1 and share large regions of homology and synteny (Figure S13), with a quarter of proteins encoded by EEC1 (24.2 %) conserved across the secondary replicons of all three species (> 70 % identity). 16 proteins were conserved with an identity above 95 % across all three replicons (Table S4). These conserved genes include *parA* and a hypothetical protein close to the replicon and two clusters of genes involved in carbohydrate metabolism, transport and regulation. Given the genetic distance of these strains, the conservation of these genes is especially notable and points to EEC1’s remarkable conservation across the genus.

The conserved regions also include the BGCs for spore pigment and, like *E. australiensis* MST-111070, the secondary replicon of *Embleya sp.* NBC 00888, also carries a BGC homologous to SapB (accession NZ_CP108785.1) (Figure S13 and S14). All replicons also share other homologous BGCs that encode the biosynthesis of specialised metabolites, most notably a homolog BGC to the siderophore peucechelin and other lanthipeptide BGCs (Figure S14). Prior to the publication of the NBC strains, five other genome assemblies of *Embleya* strains were available in the public databases (three classified as *Embleya spp.* (*Embleya hyalina* NBRC13850 GCA_003967355.1_ASM396735v1; *Embleya scabrispora* KM4927 GCA_000372745.1_ASM37274v1; and *Embleya scabrispora* NF3 GCA_002024165.1_SSNF3.0) and two wrongly classified as *Streptomyces* sp. (*Streptomyces* sp. SID3343 GCA_009865215.1_ASM986521v1; and *Streptomyces* sp. SID5474 GCA_009862895.1_ASM986289v1). These are all highly fragmented assemblies and difficult to use for analysis, apart from that of strain *E. scabrispora* NF3 whose assembly contains only 8 contigs, with one contig of over 7 Mb likely representing the nearly complete chromosome, and which does not carry the BGC for the spore pigment (instead the BGC for the spore pigment appears in a 1126 kb contig highly similar to EEC1). The *E. scabrispora* NF3 assembly also shows another 1248 kb contig with high similarity to another part of EEC1 (Figure S15) that carries BGCs for several lanthipeptides and a siderophore most similar to peuchechelin, just as EEC1 does (Figure S15). In all cases, genome alignment of *E. australiensis* MST-111070 with these five assemblies using Mauve^63^ indicates that many of the contigs of the incomplete genomes map to EEC1, providing further evidence of this replicon as a characteristic of the *Embleya* genus (Figure S16).

Also noteworthy is the fact that the *Streptomycetaceae* genus closest to *Embleya* is the rare genus *Yinghuangia*. The complete genome of *Yinghuangia* sp. ASG 101 (NCBI accession ASM2116573v1) is available. It exhibits a single replicon, the chromosome, with typical streptomycete characteristics, including the BGC for the spore pigment.

### Proposal of EEC1 as a secondary chromosome that is characteristic of the ***Embleya* genus**

The 4.2 Mb linear replicon EEC1 from *E. australiensis* MST-111070 is the largest secondary replicon described in bacteria, thus far. It is typified by a composition similar to the chromosome in terms of GC-content, codon usage and overall function, and shares the same terminal inverted repeats. EEC1 differs from the chromosome in that it contains fewer family-specific genes and is enriched in BGCs. This unique character of EEC1 makes it difficult to characterise under conventional definitions.

According to the definition introduced by Harrison et al ^36^, a chromid should meet three criteria. It should: i) utilise plasmid-like replication and maintenance systems; ii) have chromosome-like nucleotide composition and iii) carry core genes (i.e. those that are found on the chromosome of other species). Categorically, EEC1 does not fit all these criteria. In some respects, EEC1 resembles a chromid due to its chromosome-like nucleotide composition and codon usage and its genus-level distribution. However, unlike chromids, it does not appear to utilise a plasmid-like replicon or maintenance system. EEC1 appears to contain an incomplete chromosomal replicon including highly conserved homologues of ParA (> 95% identity across *Embleya* spp.) and DnaN. It also has identical TIRs to the chromosome, suggesting that its linear integrity is maintained through the same mechanism. This means that EEC1 does not meet the criteria of either a chromid or a plasmid.

Other replicons from the related genus *Streptomyces* also defy classification under this system. Most notably pSCL4 from *S. clavuligerus* ATCC 27064^5^. At 1.8 Mb, this is the largest secondary replicon from *Streptomyces* described to date. Analysis of pSCL4 under the framework provided in this study indicates that it is similar to a chromid due to its chromosomal properties (Table S5;); namely, pSCL4 encodes genes shown to be essential for chromosome stability^22^ (*tap* and *tpg*) and has codon usage similar to the chromosome. Unlike EEC1 and various chromids, it is not distributed across the genus, rather it is found uniquely in this strain. We analysed additional large *Streptomyces* plasmids, none of which exhibit the same level of similarity to the chromosome of the respective species as EEC1 does (Table S5; Figures 3).

Another important aspect is the evolutionary origin of the replicon. There are two main hypotheses for the origin of secondary replicons: the schism hypothesis, whereby the replicon is formed by a split in the chromosome, and the plasmid hypothesis whereby the replicon originates as a plasmid and attains chromosomal character over time. DiCenzo et al. took this idea further and used it as a basis for the classification of chromids and secondary chromosomes^2^. Using this approach, there is evidence to categorise EEC1 as a secondary chromosome due to its high similarity to the chromosome combined with its chromosomal-type replication and maintenance systems. However, this definition is far from accepted and there are two major problems with this system of classification. Firstly, the origin of a replicon has no bearing on its function and composition; and, secondly, it is incredibly difficult, if not impossible, to prove the evolutionary origin definitively. Genetic interchange between replicons is high and many recombinational events have been observed, such as hybridisation between secondary replicons and the chromosome^64,65^ and the interchange of terminal repeats^29^ (see also this study). Given that a defining feature of chromids is they are compositionally indistinguishable from the chromosome, it is unclear what burden of evidence would be required to definitively claim that a replicon was formed by a schism event, as opposed to a series of recombinatorial events, that would not be lost over such long evolutionary timescales. Therefore, the terms chromid, secondary chromosome, and even second chromosome, have become synonymous and are used largely interchangeably in the literature^38,39,45,65^. In summary, EEC1 shares many characteristics with chromids but lacks a plasmid-type replicon and therefore may not have originated as a plasmid. Therefore, we propose that EEC1 should be considered as a secondary chromosome but not a chromid.

EEC1 is unique and highlights the difficulty of assigning arbitrary categories to biological entities. As with any system of classification, there will always be exceptions that fall outside of the standard definitions. EEC1 and pSCL4 are two distinct examples that fall outside the established categories of chromids and megaplasmids. The study of genomic structure and secondary replicons is a developing field, particularly in *Streptomycetaceae*. While the current framework of classification provides valuable structure, it is important to recognise that secondary replicons appear to exist on a spectrum of decreasing chromosomal character and increasing evolutionary novelty.

## Conclusion

Here, we describe the genome of *Embleya australiensis* MST-111070 and its extraordinary 4.2 Mb secondary chromosome, EEC1. To our knowledge, EEC1 is the largest bacterial secondary chromosome described to date and the first secondary chromosome to be confirmed in an actinobacterial strain. Unlike other secondary chromosomes (chromids), EEC1 does not maintain itself through a plasmid-like replication system. Instead, it appears to replicate through a novel mechanism with similarities to the chromosomal replicon, suggesting it may have arisen through a separate evolutionary mechanism to other secondary chromosomes. Therefore, our research concludes that, EEC1 is a secondary chromosome but not a chromid. Most strikingly, the presence of similar secondary chromosomes for all other *Embleya* species identifies it as an essential part of the evolutionary history of the genus and a distinguishing characteristic amongst the *Streptomycetaceae*. This study underscores the importance of optical mapping in confirming the results of genome sequencing and we hope the discovery of EEC1 will encourage further, in-depth characterisation of extraordinary bacterial replicons.

## Materials and Methods

### Strains and culture conditions

*E. australiensis* MST-111070 was isolated from arid soil collected in South Australia in 1996. Cultivation was performed on SFM agar (20 g/L Soya Flour, 20 g/L Mannitol)^10^ for sporulation; spore stocks were prepared following established methods^10^.

### DNA extraction and quality control

High molecular weight DNA samples were extracted from early-stationary-phase liquid cultures in 1:1 TSB:YEME medium^10^, following the salting-out protocol^10^. Sample quality was first assessed by standard agarose-gel electrophoresis and pulse-field gel electrophoresis, Quantification and further quality control was performed with NanoDrop and Qubit (Thermo Fisher Scientific) following manufacturers’ instructions, before handing the samples to sequencing providers.

### Genome sequencing and assembly

PacBio SMRT sequencing and assembly (Pacific Biosciences of California, Inc) was commissioned to the Earlham Institute (Norwich Research Park, Norwich, NR4 7UZ, United Kingdom). Sequencing was performed with C4-P6 chemistry on three SMRT cells with a RSII sequencer; the data were processed and assembled with HGAP.2 and HGAP.3. Illumina sequencing and assembly was commissioned to the DNA sequencing facility in the Department of Biochemistry, University of Cambridge (Cambridge, CB2 1GA, UK). Details given by the sequencing provider: a shotgun library with average insert size of 550 bp was prepared using the Illumina TruSeq PCRfree/Nano library preparation kit; a second shotgun library with average insert size of 4 kb was prepared using Illumina Nextera Mate Pair library preparation kit; both libraries were sequenced using V2 Illumina sequencing chemistry and run on a MiSeq instrument (2 × 250-bp paired-end sequencing). Sequence data was processed and assembled with a custom pipeline developed at the facility (Dr Markiyan Samborskyy, unpublished data) and as previously reported^21^.

### Optical mapping

Optical mapping was performed with Bionano Irys (Bionano Genomics, San Diego, USA) technology. Bacterial cells were prepared from a liquid culture following supplier’s instructions; DNA-extraction, labelling and data collection including processing of images to extract BNX files with molecules information, was commissioned to the Earlham Institute (Norwich Research Park, Norwich, NR4 7UZ, United Kingdom). We obtained BNX files with Bionano molecules information for a high-resolution map of the distribution of the recognition site for nickase Nt.BspQI (GCTCTTC). The BNX files were processed with Bionano’s software IrysView Genomic Analysis Viewer, version 2.5.1.29842, with versions r5134 of PipelineCL.py, r5122 of RefAligner.cpp, r5122 of Assembler.cpp, and r5146 of Hybrid Scaffold; all running on Microsoft Windows 10 64 bits with Python 2.7.8. The software was used following the guidelines in the Bionano document “IrysView® v2.5.1 Software Training Guide. Document Number: 30035 Document Revision: G, 2016”. The result of Molecule Quality Report (alignment over reference sequence) was interpreted according to the document “Guidelines for Interpreting the Bionano Molecule Quality Report. Document number 30175, Rev A”. Our analysis strongly benefited from invaluable advice provided by BioNano Technical Support. A detailed account of the methodology is given in Supporting Information.

### Sequence data analysis

General visualisation, analysis, and manipulation of DNA sequence data was performed with computer programs Artemis^66^, Artemis Comparison Tool^67^, and NotePad++ (http://notepad-plus-plus.org/). Alignments and assembly of sequences was performed with Staden Package^68,69^. Similarity searches were performed with BLAST+^70^at the NCBI web server (http://www.ncbi.nlm.nih.gov/blast/), or on a standalone computer with prfectBLAST 2.0^71^. Annotation of gene function and genetic features was performed with antiSMASH (version 7 with “relaxed strictness” and all options enabled)^72^ and RAST (Rapid Annotation using Subsystem Technology)^73^ using Prodigal^74^ for CDS calling, and with frameshift fixing, and metabolic model building options. Mauve^63^ was used for full genome comparisons and contigs reordering. BUSCO^46^ was used to analyse the presence of orthologous genes per taxonomic group (run last time on 19th March 2023, version BUSCO v5.4.4). Dot plots were generated using dotplotter v1.0.0^75^.

### Taxonomic analysis

TYGS^33^ and GGDC^76^ were used for genome-based and 16S rDNA-based taxonomical studies (with Extended Maximum Likelihood and Maximum Parsimony 16S rRNA analysis, which includes type strains for which the genome sequence is not available). All taxonomical investigations, as well as genome alignments with Mauve^63^ were last repeated during May and June 2023 prior to the finalisation of this manuscript, as to reflect the most current state of genome and 16S rDNA sequence availability in the databases. The 11-locus tree was produced by running getphylo^35^ on 515 complete genomes accessed from the NCBI (21/02/24) and visualised using iTOL^77^.

### Functional and compositional analysis

GC content and GC skew was calculated with a custom python package (gcskewer) developed for this project (https://github.com/drboothtj/gcskewer). Codon usage and codon usage correlation was calculated using a python package (codoniser) developed for this project. codoniser is open-source and is available on the Python Package Index (PyPI) and GitHub (https://github.com/drboothtj/codoniser). Replicon function was estimated using COG categories as annotated in the EggNOG 5.0^47^ database and implemented in eggnog-mapper v2^48^ with the taxonomic scope set to ‘Actinobacteria’. COG category correlation was calculated using a python package (egger) developed for this project and available on PyPI and GitHub (https://github.com/drboothtj/egger). The rank-specificity of genes was calculated using diamond blastp searches against the predicted proteomes of the complete genomes used in the taxonomic analyses above. Specificity scores were calculated for each protein coding sequence using the following equation:

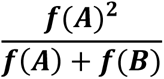

Where, 𝒇(𝑨) is the frequency of the gene occurring on a genome in group A and 𝒇(𝑩) is the frequency of the gene occurring on a genome in group B. For strain specificity, group A contained only MST-111070 and group B included all other taxa. For genus-specificity, group A contained all the *Embleya* sequences and group B included all other taxa. Finally, for family-specificity, group A contained all the *Embleya*, *Streptomyces, Kitosatospora* and *Yinghuangia* sequences and group B included all other taxa.

## Acknowledgements

This work was supported by Fellowship funding from the Daphne Jackson Trust and the Biotechnology and Biological Sciences Research Council (BBSRC) to S.D-R., by BBSRC via Responsive Mode Grant BB/P021506/1 to B.W., by BBSRC Institute Strategic Programme Grants BBS/E/J/000PR9790 and BB/X01097X/1 to the John Innes Centre (JIC), by BBSRC Doctoral Training Program Grant BB/M011216/1 to T.J.B., and by Novo Nordisk Foundation Grant NNF22OC0078997 to T.J.B. We acknowledge the Earlham Institute (Norwich, UK) for PacBio sequencing, Bionano optical mapping, and initial assemblies. We acknowledge the University of Cambridge (Department of Biochemistry) DNA sequencing facility for Illumina sequencing and initial assemblies. We thank Professor Mervyn Bibb (John Innes Centre) for helpful discussions and invaluable advice during the drafting of this manuscript.

## Author contributions

**J.P.G-E.:** methodology, validation, formal analysis, investigation, data curation, writing (original draft, review & editing), visualization. **S.D-R.:** methodology, validation, investigation. **D.B.:** methodology, validation, investigation. **E.L.:** conceptualization, resources, writing (review & editing), project administration. **B.W.:** conceptualization, methodology, validation, resources, writing (review & editing), supervision, project administration, funding acquisition. **T.J.B.:** methodology, validation, formal analysis, investigation, data curation, writing (original draft, review & editing), visualization, funding acquisition.

## References

1. Kinashi, H. & Shimaji-Murayama, M. Physical characterization of SCP1, a giant linear plasmid from Streptomyces coelicolor. J Bacteriol 173, 1523 (1991).

2. diCenzo, G. C. & Finan, T. M. The Divided Bacterial Genome: Structure, Function, and Evolution. Microbiol Mol Biol Rev 81, e00019–17 (2017).

3. Kinashi, H. Giant linear plasmids in Streptomyces: a treasure trove of antibiotic biosynthetic clusters. J Antibiot (Tokyo) 64, 19–25 (2011).

4. Li, P., Zhang, J., Deng, Z., Gao, F. & Ou, H.-Y. Identification and characterization of a central replication origin of the mega-plasmid pSCATT of Streptomyces cattleya. Microbiol Res 257, 126975 (2022).

5. Medema, M. H. et al. The sequence of a 1.8-mb bacterial linear plasmid reveals a rich evolutionary reservoir of secondary metabolic pathways. Genome Biol Evol 2, 212–224 (2010).

6. Letek, M. et al. Evolution of the Rhodococcus equi vap Pathogenicity Island Seen through Comparison of Host-Associated vapA and vapB Virulence Plasmids. J Bacteriol 190, 5797 (2008).

7. Francis, I. et al. pFiD188, the Linear Virulence Plasmid of Rhodococcus fascians D188. / 637 MPMI 25, 637–647 (2012).

8. Alvarez-Narvaez, S., Giguere, S., Berghaus, L. J., Dailey, C. & Vazquez-Boland, J. A. Horizontal Spread of Rhodococcus equi Macrolide Resistance Plasmid pRErm46 across Environmental Actinobacteria. Appl Environ Microbiol 86, e00108–20 (2020).

9. Parra, J. et al. Antibiotics from rare actinomycetes, beyond the genus Streptomyces. Curr Opin Microbiol 76, 102385 (2023).

10. Kieser, T. et al. Practical Streptomyces Genetics. (John Innes Foundation, 2000).

11. Flärdh, K. & Buttner, M. J. Streptomyces morphogenetics: dissecting differentiation in a filamentous bacterium. Nat Rev Microbiol 7, 36–49 (2009).

12. Nouioui, I. et al. Genome-based taxonomic classification of the phylum actinobacteria. Front Microbiol 9, 355158 (2018).

13. Awano, Y. et al. Emblestatin: a new peptide antibiotic from Embleya scabrispora K20-0267. J Antibiot (Tokyo) 76, 592–597 (2023).

14. Parte, A. C., Carbasse, J. S., Meier-Kolthoff, J. P., Reimer, L. C. & Göker, M. List of Prokaryotic names with Standing in Nomenclature (LPSN) moves to the DSMZ. Int J Syst Evol Microbiol 70, 5607–5612 (2020).

15. Ōmura, S., Nakagawa, A., Shibata, K. & Sano, H. The structure of hitachimycin, a novel macrocyclic lactam involving β-phenylalanine. Tetrahedron Lett 23, 4713–4716 (1982).

16. Kudo, F. et al. Genome Mining of the Hitachimycin Biosynthetic Gene Cluster: Involvement of a Phenylalanine-2,3-aminomutase in Biosynthesis. ChemBioChem 16, 909–914 (2015).

17. Hiramatsu, K. et al. Curing bacteria of antibiotic resistance: reverse antibiotics, a novel class of antibiotics in nature. Int J Antimicrob Agents 39, 478–485 (2012).

18. Estévez, M. R., Myronovskyi, M., Gummerlich, N., Nadmid, S. & Luzhetskyy, A. Heterologous Expression of the Nybomycin Gene Cluster from the Marine Strain Streptomyces albus subsp. chlorinus NRRL B-24108. Mar Drugs 16, 435 (2018).

19. Rodríguez-Peña, K. et al. Bioinformatic comparison of three Embleya species and description of steffimycins production by Embleya sp. NF3. 1, 3 (1915).

20. Jørgensen, T. S. et al. A treasure trove of 1034 actinomycete genomes. Nucleic Acids Res 52, 7487– 7503 (2024).

21. Oliveira, L. G. de, Sigrist, R., Paulo, B. S. & Samborskyy, M. Whole-Genome Sequence of the Endophytic Streptomyces sp. Strain CBMAI 2042, Isolated from Citrus sinensis. Microbiol Resour Announc 8, e01426–18 (2019).

22. Gomez-Escribano, J. P. et al. Genome editing reveals that pSCL4 is required for chromosome linearity in Streptomyces clavuligerus. Microb Genom 7, 000669 (2021).

23. Gomez-Escribano, J. P., Alt, S. & Bibb, M. J. Next Generation Sequencing of Actinobacteria for the Discovery of Novel Natural Products. Mar Drugs 14, 78 (2016).

24. Chen, C. W., Huang, C. H., Lee, H. H., Tsai, H. H. & Kirby, R. Once the circle has been broken: dynamics and evolution of Streptomyces chromosomes. Trends Genet 18, 522–529 (2002).

25. Weaver, D. et al. Genome plasticity in Streptomyces: identification of 1 Mb TIRs in the S. coelicolor A3(2) chromosome. Mol Microbiol 51, 1535–1550 (2004).

26. Lezhava, A. et al. Physical map of the linear chromosome of Streptomyces griseus. J Bacteriol 177, 6492 (1995).

27. Fischer, G., Decaris, B. & Leblond, P. Occurrence of deletions, associated with genetic instability in Streptomyces ambofaciens, is independent of the linearity of the chromosomal DNA. J Bacteriol 179, 4553 (1997).

28. Volff, J. N., Viell, P. & Altenbuchner, J. Artificial circularization of the chromosome with concomitant deletion of its terminal inverted repeats enhances genetic instability and genome rearrangement in Streptomyces lividans. Molecular and General Genetics 253, 753–760 (1997).

29. Chen, C. W., Huang, C. H., Lee, H. H., Tsai, H. H. & Kirby, R. Once the circle has been broken: Dynamics and evolution of Streptomyces chromosomes. Trends in Genetics 18, 522–529 (2002).

30. Meng, Y. et al. Genome sequence assembly algorithms and misassembly identification methods. Mol Biol Rep 49, 11133–11148 (2022).

31. Leinonen, M. & Salmela, L. Optical map guided genome assembly. BMC Bioinformatics 21, 1–19 (2020).

32. Yuan, Y., Chung, C. Y.-L. & Chan, T.-F. Advances in optical mapping for genomic research. Comput Struct Biotechnol J 18, 2051–2062 (2020).

33. Meier-Kolthoff, J. P. & Göker, M. TYGS is an automated high-throughput platform for state-of-the-art genome-based taxonomy. Nat Commun 10, 1–10 (2019).

34. Meier-Kolthoff, J. P., Carbasse, J. S., Peinado-Olarte, R. L. & Göker, M. TYGS and LPSN: a database tandem for fast and reliable genome-based classification and nomenclature of prokaryotes. Nucleic Acids Res 50, D801–D807 (2022).

35. Booth, T. J., Shaw, S., Cruz-Morales, P. & Weber, T. getphylo: rapid and automatic generation of multi-locus phylogenetic trees. BMC Bioinform 26, 1–11 (2025).

36. Harrison, P. W., Lower, R. P. J., Kim, N. K. D. & Young, J. P. W. Introducing the bacterial ‘chromid’: not a chromosome, not a plasmid. Trends Microbiol 18, 141–148 (2010).

37. Slater, S. C. et al. Genome sequences of three agrobacterium biovars help elucidate the evolution of multichromosome genomes in bacteria. J Bacteriol 191, 2501–2511 (2009).

38. Bochkareva, O. O., Moroz, E. V., Davydov, I. I. & Gelfand, M. S. Genome rearrangements and selection in multi-chromosome bacteria Burkholderia spp. BMC Genomics 19, 1–17 (2018).

39. Dicenzo, G. C., Mengoni, A. & Perrin, E. Chromids Aid Genome Expansion and Functional Diversification in the Family Burkholderiaceae. Mol Biol Evol 36, 562–574 (2019).

40. Xie, B. Bin et al. Evolutionary trajectory of the replication mode of bacterial replicons. mBio 12, 1– 15 (2021).

41. Jiang, W. et al. Mutational features of chromids and chromosomes in Pseudoalteromonas provide new insights into the evolution of secondary replicons. Microbiol Spectr 13, (2025).

42. Suwanto, A. & Kaplan, S. Physical and genetic mapping of the Rhodobacter sphaeroides 2.4.1 genome: presence of two unique circular chromosomes. J Bacteriol 171, 5850–5859 (1989).

43. Okada, K., Iida, T., Kita-Tsukamoto, K. & Honda, T. Vibrios Commonly Possess Two Chromosomes. J Bacteriol 187, 752 (2005).

44. Riccardi, C. et al. Independent origins and evolution of the secondary replicons of the class Gammaproteobacteria. Microb Genom 9, 001025 (2023).

45. Fournes, F. et al. The coordinated replication of Vibrio cholerae’s two chromosomes required the acquisition of a unique domain by the RctB initiator. Nucleic Acids Res 49, 11119 (2021).

46. Manni, M., Berkeley, M. R., Seppey, M. & Zdobnov, E. M. BUSCO: Assessing Genomic Data Quality and Beyond. Curr Protoc 1, e323 (2021).

47. Huerta-Cepas, J. et al. eggNOG 5.0: a hierarchical, functionally and phylogenetically annotated orthology resource based on 5090 organisms and 2502 viruses. Nucleic Acids Res 47, D309–D314 (2019).

48. Cantalapiedra, C. P., Hern̗andez-Plaza, A., Letunic, I., Bork, P. & Huerta-Cepas, J. eggNOG-mapper v2: Functional Annotation, Orthology Assignments, and Domain Prediction at the Metagenomic Scale. Mol Biol Evol 38, 5825–5829 (2021).

49. Gilchrist, C. L. M. et al. cblaster: a remote search tool for rapid identification and visualisation of homologous gene clusters. Bioinform Adv 1, 1–10 (2021).

50. Blin, K. et al. antiSMASH 7.0: new and improved predictions for detection, regulation, chemical structures and visualisation. Nucleic Acids Res 51, W46–W50 (2023).

51. Du, Y. et al. Identification and characterization of the biosynthetic gene cluster of polyoxypeptin A, a potent apoptosis inducer. BMC Microbiol 14, 30 (2014).

52. Hu, Z. & Reid, R. Biosynthetic gene cluster for leptomycins. (2007).

53. Willey, J., Santamaria, R., Guijarro, J., Geistlich, M. & Losick, R. Extracellular complementation of a developmental mutation implicates a small sporulation protein in aerial mycelium formation by S. coelicolor. Cell 65, 641–650 (1991).

54. Kodani, S. et al. The SapB morphogen is a lantibiotic-like peptide derived from the product of the developmental gene ramS in Streptomyces coelicolor. Proc Natl Acad Sci U S A 101, 11448–11453 (2004).

55. Chang, P.-C. & Cohen, S. N. Bidirectional replication from an internal origin in a linear streptomyces plasmid. Science 265, 952–954 (1994).

56. Redenbach, M., Bibb, M., Gust, B., Seitz, B. & Spychaj, A. The Linear Plasmid SCP1 of Streptomyces coelicolor A3(2) Possesses a Centrally Located Replication Origin and Shows Significant Homology to the Transposon Tn4811. Plasmid 42, 174–185 (1999).

57. Mochizuki, S. et al. The large linear plasmid pSLA2-L of Streptomyces rochei has an unusually condensed gene organization for secondary metabolism. Mol Microbiol 48, 1501–1510 (2003).

58. Musialowski, M. S. et al. Functional evidence that the principal DNA replication origin of the Streptomyces coelicolor chromosome is close to the dnaA-gyrB region. J Bacteriol 176, 5123–5125 (1994).

59. Calcutt, M. J. Gene organization in the dnaA-gyrA region of the Streptomyces coelicolor chromosome. Gene 151, 23–28 (1994).

60. Smulczyk-Krawczyszyn, A., et al. Cluster of DnaA Boxes Involved in Regulation of Streptomyces Chromosome Replication: from In Silico to In Vivo Studies. J Bacteriol 188, 6184 (2006).

61. Bignell, C. & Thomas, C. M. The bacterial ParA-ParB partitioning proteins. J Biotechnol 91, 1–34 (2001).

62. Bentley, S. D. et al. SCP1, a 356 023 bp linear plasmid adapted to the ecology and developmental biology of its host, Streptomyces coelicolor A3(2). Mol Microbiol 51, 1615–1628 (2004).

63. Rissman, A. I. et al. Reordering contigs of draft genomes using the Mauve aligner. Bioinformatics 25, 2071–2073 (2009).

64. Yamasaki, M. & Kinashi, H. Two chimeric chromosomes of Streptomyces coelicolor A3 (2) generated by single crossover of the wild-type chromosome and linear plasmid SCP1. J Bacteriol 186, 6553– 6559 (2004).

65. Mori, J. F. & Kanaly, R. A. Natural Chromosome-Chromid Fusion across rRNA Operons in a Burkholderiaceae Bacterium. Microbiol Spectr 10, (2022).

66. Rutherford, K., et al. Artemis: sequence visualization and annotation. Bioinformatics 16, 944–945 (2000).

67. Carver, T. J. et al. ACT: the Artemis Comparison Tool. Bioinformatics 21, 3422–3423 (2005).

68. Staden, R., Beal, K. F. & Bonfield, J. K. The Staden package, 1998. Methods Mol Biol 132, 115–130 (2000).

69. Bonfield, J. K. & Whitwham, A. Gap5—editing the billion fragment sequence assembly. Bioinformatics 26, 1699 (2010).

70. Altschul, S. F. et al. Gapped BLAST and PSI-BLAST: a new generation of protein database search programs. Nucleic Acids Res 25, 3389–3402 (1997).

71. Santiago-Sotelo, P. & Ramirez-Prado, J. H. prfectBLAST: a platform-independent portable front end for the command terminal BLAST+ stand-alone suite. Biotechniques 53, 299–300 (2012).

72. Blin, K. et al. antiSMASH 7.0: new and improved predictions for detection, regulation, chemical structures and visualisation. Nucleic Acids Res 51, W46–W50 (2023).

73. Aziz, R. K. et al. The RAST Server: Rapid Annotations using Subsystems Technology. BMC Genomics 9, 75 (2008).

74. Hyatt, D. et al. Prodigal: prokaryotic gene recognition and translation initiation site identification. BMC Bioinformatics 11, 119 (2010).

75. Mohite, O. S. et al. Pangenome mining of the Streptomyces genus redefines species’ biosynthetic potential. Genome Biology 2025 26:1 26, 1–20 (2025).

76. Meier-Kolthoff, J. P., Auch, A. F., Klenk, H. P. & Göker, M. Genome sequence-based species delimitation with confidence intervals and improved distance functions. BMC Bioinformatics 14, 1– 14 (2013).

77. Letunic, I. & Bork, P. Interactive tree of life (iTOL) v3: an online tool for the display and annotation of phylogenetic and other trees. Nucleic Acids Res 44, W242–5 (2016).

78. Price, M. N., Dehal, P. S. & Arkin, A. P. FastTree 2 - Approximately maximum-likelihood trees for large alignments. PLoS One 5, e9490 (2010).

79. Gilchrist, C. L. M. & Chooi, Y.-H. clinker & clustermap.js: automatic generation of gene cluster comparison figures. Bioinform 37, 2473–2475 (2021).

80. Mistry, J. et al. Pfam: The protein families database in 2021. Nucleic Acids Res 49, D412–D419 (2021).

