## Supplementary Information for "Evidence supporting the first secondary chromosome in actinobacteria as a hallmark of the *Embleya* genus"

### EXTENDED METHODS DESCRIPTION

#### Sequence data analysis

General visualisation, analysis, and manipulation of DNA sequence data was performed with computer programs Artemis[1], Artemis Comparison Tool[2], and NotePad++ (http://notepad-plus-plus.org/). Alignments and assembly of sequences was performed with Staden Package[3, 4]. Similarity searches were performed with BLAST+ [5] at the NCBI web server (http://www.ncbi.nlm.nih.gov/blast/), or on a standalone computer with prfectBLAST 2.0[6]. Annotation of gene function and genetic features was performed with antiSMASH (version 7 with “relaxed strictness” and all options enabled)[7] and RAST (Rapid Annotation using Subsystem Technology)[8, 9] using Prodigal[10] for CDS calling, and with frame-shift fixing, and metabolic model building options. Mauve[11] was used for full genome comparisons and contigs reordering. Busco[12] was used to analyse the presence of orthologous genes per taxonomic group (version BUSCO v5.4.4 with hmmsearch 3.1, bbtools 39.01, prodigal 2.6.3, with databases actinobacteria_class_odb10; actinobacteria_phylum_odb10; bacteria_odb10; streptomycetales_odb10). TYGS[13] and GGDC[14, 15] were used for genome-based and 16S rDNA-based taxonomical studies (with Extended Maximum Likelihood and Maximum Parsimony 16S rRNA analysis, which includes type strains for which the genome sequence is not available).

#### Bionano data analysis

The BNX files with Bionano molecules information were processed with Bionano’s software IrysView Genomic Analysis Viewer, version 2.5.1.29842, with versions r5134 of PipelineCL.py, r5122 of RefAligner.cpp, r5122 of Assembler.cpp, and r5146 of Hybrid Scaffold; all running on Microsoft Windows 10 64 bits with Python 2.7.8. IrysView was used for alignment of Bionano molecules over the consensus genome sequence and visualisation, following the guidelines in the Bionano document “IrysView® v2.5.1 Software Training Guide. Document Number: 30035 Document Revision: G, 2016”; the result of Molecule Quality Report (alignment over reference sequence) was interpreted according to the document “Guidelines for Interpreting the Bionano Molecule Quality Report. Document number30175, Rev A” and following advice from BioNano Technical Support.

During import of BNX files into IrysView, molecules were filtered for a minimum of 100 kb length with Dynamic Threshold label SNR.

Data for all scans were merged with default options, resulting in a total of 229509 molecules accounting for 37288.4 Mb, with a molecule N50 of 158.5 Kb.

##### Molecules-to-Reference Alignment

**Molecule Quality Report (MQR)** aligns the Bionano molecules to a reference sequence, being a good indicator of the accuracy of the DNA-sequence assembly. MQR was performed without molecule subsampling, 3 iterations, and the following molecule-to-reference RefAligner options:

```
-nosplit 2 -BestRef 1 -biaswt 0 -Mfast 0 -FP 1.5 -FN 0.15 -sf 0.2 -sd 0.0 -A 5 -
outlier 1e-3 -outlierMax 40 -endoutlier 1e-4 -S -1000 -sr 0.03 -se 0.2 -MaxSF
0.25 -MaxSE 0.5 -resbias 4 64 -maxmem 64 -M 3 3 -minlen 150 -T 1e-8 -maxthreads
32 -hashgen 5 3 2.4 1.5 0.05 5.0 1 1 3 -hash -hashdelta 10 -hashoffset 1 -
hashmaxmem 64 -insertThreads 4 -maptype 0 -PVres 2 -PVendoutlier -AlignRes 2.0 -
rres 0.9 -resEstimate -ScanScaling 2 -RepeatMask 5 0.01 -RepeatRec 0.7 0.6 1.4 -
maxEnd 50 -usecolor 1 -stdout –stderr
```

The only parameter changed in relation with the default options was the T option (the P-value cut-off) from 1e-11 (set for human genome) to 1e-8; this is still one order of magnitude more stringent than the suggested for small genomes, 1e-7 for *Escherichia coli* and 5 Mbp genome size (see Bionano's documentation "Guidelines for Interpreting the Bionano Molecule Quality Report. Document number30175, Rev A"

As reference DNA sequence we used version 4 of the assembly (PacBio HGAP.3, TIRs extended with Illumina, putative circular plasmid sequence starting with *parB* stop codon).

Despite the Map Rate was 42.7% (Molecules showing high similarity to reference as percentage of 'N Molecules') which is below the expected range (60-80%), the relevant statistic indicators for the quality of mapping were well within the range established by Bionano, at FP 0.48/100kbp, FP 4.4%, and FN 18.9% (limits stated in the Guidelines are FP (/100kbp) < 1.7; FP (%) < 15%; FN (%) < 21%; "FP (/100kb)" means density of molecule labels absent in the reference map (relative to reference labels); "FP (%)" means percentage of molecule labels absent in the reference map (relative to reference labels); "FN (%)" means percentage of reference labels absent in the aligned molecules (relative to reference labels); see Bionano's document number 30175 for details)

Therefore, the statistics alone indicate that the Bionano data align well with the reference sequence, supporting the correctness of our assembly version 4. Beyond the mere statistics, detailed study of the alignment reveals the excellent distribution of Bionano molecules along the DNA assembly (see Figure S2).

**De novo assembly of Bionano molecules**

Assembly was performed with the merged filtered data as before, using the settings for small size genomes (OptArgs: optArguments_small) and without reference.

The result was 8 contigs spanning a total of 14.359 Mb, with N50 of 4.33 Mb and an average coverage of 106.3 molecules:

| Contig | Map ID | Size |
|---|---|---|
| 1 | 1 | 4.330 Mb |
| 2 | 2 | 4.363 Mb |
| 3 | 4 | 2.022 Mb |
| 4 | 11 | 556.986 kb |
| 5 | 17 | 578.444 kb |
| 6 | 125 | 551.393 kb |
| 7 | 231 | 1.614 Mb |
| 8 | 1207 | 343.984 kb |

**Alignment of BioNano De novo assembly to DNA-sequence assembly**

The de-novo assembled bionano contigs were mapped and aligned to the DNA sequence assembly version 4 provided as reference, with the following options:

```
-maxthreads 32 -output-veto-filter _intervals.txt$ -res 2.9 -FP 0.6 -FN 0.06 -sf 0.20 -sd 0.0 -sr 0.01 -extend 1 -outlier 0.0001 -endoutlier 0.001 -PVendoutlier -deltaX 12 -deltaY 12 -xmapchim 12 -hashgen 5 7 2.4 1.5 0.05 5.0 1 1 1 -hash -hashdelta 50 -mres 1e-3 -hashMultiMatch 100 -insertThreads 4 -nosplit 2 -biaswt 0 -T 1e-8 -S -1000 -indel -PVres 2 -rres 0.9 -MaxSE 0.5 -HSDrange 1.0 -outlierBC -xmapUnique 12 -AlignRes 2. -outlierExtend 12 24 -Kmax 12 -f -maxmem 128 -BestRef 1 -stdout -stderr
```

### SUPPLEMENTARY TABLES

**Table S1. Loci selected for phylogenetic analysis by getphylo. The locus ID refers to the locus tag from the the seed genome *Streptomyces* sp. LX-29.**

| Locus ID | Alignment Length | Informative Sites | Description |
|---|---|---|---|
| LRS74_RS08715 | 479 | 276 | pseudouridine synthase B (TruB) |
| LRS74_RS08740 | 435 | 212 | transcription termination/antitermination protein NusA |
| LRS74_RS10140 | 494 | 209 | F0F1 ATP synthase subunit beta |
| LRS74_RS13285 | 379 | 197 | tRNA threonylcarbamoyladenosine biosynthesis protein TsaB |
| LRS74_RS13530 | 356 | 127 | small ribosomal subunit protein uS3 |
| LRS74_RS13545 | 279 | 81 | large ribosomal subunit protein uL2 |
| LRS74_RS13620 | 243 | 110 | 50S ribosomal protein L1 |
| LRS74_RS17975 | 207 | 104 | DUF47 domain-containing protein |
| LRS74_RS20580 | 361 | 241 | ribosome hibernation promotion factor (HPF) |
| LRS74_RS23160 | 320 | 191 | nicotinate-nucleotide adenylyltransferase |
| LRS74_RS24320 | 315 | 203 | zinc ribbon domain-containing protein |

**Table S2. Identification of genes associated with chromosomal or plasmid origins of replication.** The results show the percentage identity of hits from *Embleya australiensis* MST-111070 to corresponding genes from *Streptomyces clavuligerus* ATCC27064 from Diamond[16] BlastP searches. Blank cells indicate no hit.

| replicon | Chromosome | | | | pSCL4 | | | |
|---|---|---|---|---|---|---|---|---|
| | *dnaA* | *dnaN* | *parA* | *parB* | *rep1* | *rep2* | *parA* | *parB* |
| Chromosome | 55.9 | 75.9 | 82.9 | 57.1 | - | - | 34.1 | 30.1 |
| EEC1 | - | 43.3 | 42.1 | - | - | - | - | - |
| EEC2 | - | - | 34.6 | 36.9 | - | - | 38.9 | - |
| EEC3 | - | - | 44.6 | - | 73.4 | 74.2 | - | 47.3 |

**Table S3. Curated antiSMASH output for the genome of *Embleya australiensis* MST-111070.**

| Region/ BGC[1] | Type[2] | From nt[3] | To nt[3] | Most similar known BGC[4] |
|---|---|---|---|---|
| Chr-1 | T1PKS,linaridin | 89,951 | 138,901 | *Legonaridin (BGC0001188)* |
| Chr-2 | terpene | 150,625 | 171,617 | *None* |
| Chr-3 | CDPS | 171,621 | 192,325 | *None* |
| Chr-4 | NRPS-like,NRPS,T1PKS | 195,790 | 343,903 | *Incednine (BGC0000078)* |
| Chr-5 | thiopeptide,LAP,T1PKS | 349,551 | 414,513 | *Paromomycin (BGC0000712)* |
| Chr-6 | terpene | 882,536 | 903,615 | *Dudomycin A (BGC0002359)* |
| Chr-7 | indole,NRPS,T1PKS | 980,466 | 1,101,259 | **Antibiotic L 156602[5]** |
| Chr-8 | NRP-metallophore,NRPS | 1,177,888 | 1,235,978 | Griseobactin (BGC0000368) |
| Chr-9 | T3PKS | 1,266,421 | 1,307,485 | Alkylresorcinol (BGC0000282) |
| Chr-10 | other | 1,336,893 | 1,377,618 | *Himastatin (BGC0001117)* |
| Chr-11 | hydrogen-cyanide | 1,439,526 | 1,452,331 | *Aborycin (BGC0002285)* |
| Chr-12 | NRPS,HR-T2PKS | 4,277,131 | 4,345,962 | Ishigamide (BGC0001623) |
| Chr-13 | NAPAA,T1PKS,hglE-KS | 5,295,805 | 5,365,248 | *Hexacosalactone A (BGC0002497)* |
| Chr-14 | lanthipeptide-class-iv | 5,425,093 | 5,448,404 | *Labyrinthopeptin A1-3 (BGC0000519)* |
| Chr-15 | NI-siderophore | 5,479,577 | 5,513,301 | *Schizokinen (BGC0002683)* |
| Chr-16 | NI-siderophore | 5,640,743 | 5,670,680 | *None* |
| Chr-17 | RiPP-like | 5,701,274 | 5,712,086 | *None* |
| Chr-18 | terpene,betalactone | 5,803,329 | 5,857,721 | Hopene (BGC0000663) |
| Chr-19 | NRPS-like,betalactone | 5,940,563 | 5,984,102 | **Nybomycin (BGC0001965)** |
| Chr-20 | T1PKS | 6,126,397 | 6,199,439 | *Neoabyssomicin/Abyssomicin (BGC0001694)* |
| Chr-21 | terpene | 6,559,542 | 6,581,863 | *Polyoxypeptin (BGC0001036)* |
| Chr-22 | RiPP-like | 6,595,549 | 6,606,349 | *Persiamycin A (BGC0002045)* |
| Chr-23 | terpene | 6,778,221 | 6,799,270 | *None* |
| Chr-24 | T1PKS | 6,905,975 | 7,081,035 | **Leptomycin/Kazusamycin[6]** |
| EEC1-1 | PKS-like | 294,513 | 335,535 | *Rustmicin (BGC0000065)* |
| EEC1-2 | NRPS,NRPS-like | 524,754 | 578,809 | *WS9326 (BGC0001297)* |
| EEC1-3 | NRPS,NRPS-like | 608,729 | 676,086 | Himastatin (BGC0001117) |
| EEC1-4 | NRPS,betalactone | 683,228 | 753,983 | *Ohmyungsamycin A-B (BGC0002293)* |
| EEC1-5 | terpene,NRPS, aminocoumarin,HR-T2PKS | 971,183 | 1,044,988 | *Acyldepsipeptide 1 (BGC0001967)* |

| | | | | |
|---|---|---|---|---|
| EEC1-6 | RiPP-like | 1,049,383 | 1,061,281 | *None* |
| EEC1-7 | RRE-containing, butyrolactone | 1,064,960 | 1,091,725 | *None* |
| EEC1-8 | NAPAA | 1,336,612 | 1,370,679 | *γ-poly-L-2,4-diaminobutyric acid (BGC0002535)* |
| EEC1-9 | hydrogen-cyanide | 1,462,193 | 1,475,159 | *None* |
| EEC1-10 | NRPS | 1,550,546 | 1,593,887 | *Triacsin C (BGC0001983)* |
| EEC1-11 | lanthipeptide-class-iv | 1,619,382 | 1,641,973 | *Ulleungmycin (BGC0001814)* |
| EEC1-12 | lanthipeptide-class-iii | 1,828,683 | 1,851,442 | SapB homologue[7] |
| EEC1-13 | NRPS | 2,014,404 | 2,064,860 | |
| EEC1-14 | terpene | 2,106,657 | 2,127,820 | *None* |
| EEC1-15 | terpene | 2,240,391 | 2,261,305 | *None* |
| EEC1-16 | NRPS,T1PKS | 2,426,285 | 2,492,967 | *ECO-0501 (BGC0002098)* |
| EEC1-17 | NRPS,lanthipeptide-class-ii | 2,579,389 | 2,634,742 | *None* |
| EEC1-18 | NI-siderophore,NRPS | 2,641,507 | 2,731,521 | *Peucechelin (BGC0002466)* |
| EEC1-19 | T2PKS | 2,792,990 | 2,865,499 | Spore Pigment (BGC0000271)[8] |
| EEC1-20 | T3PKS | 2,869,003 | 2,910,154 | *R1128 (BGC0000261)* |
| EEC1-21 | T1PKS,HR-T2PKS, arylpolyene,NRPS-like | 3,295,026 | 3,442,703 | *Akaeolide (BGC0001199)* |
| EEC1-22 | hglE-KS | 3,522,775 | 3,573,744 | *None* |
| EEC1-23 | lassopeptide | 3,725,618 | 3,748,424 | *None* |
| EEC1-24 | terpene | 3,796,308 | 3,817,516 | *Ebelactone (BGC0001580)* |
| EEC1-25 | NRPS-like | 3,890,578 | 3,933,670 | *None* |
| EEC1-26 | NRPS-like | 4,013,519 | 4,071,452 | *Ecumicin (BGC0001582)* |
| EEC2-1 | lanthipeptide-class-i | 120,682 | 145,886 | *None* |
| EEC3-1 | ranthipeptide | 36,794 | 75,454 | *Desulfoclethramycin/Clethramycin (BGC0002498)* |

[1]antiSMASH "regions" might include one or more BGCs; "Chr" states for "chromosome". [2]antiSMASH annotation. [3]Beguining and end positions of the "region" as annotated by antiSMASH. [4]Know BGC with which the "region" contains highest proportion of similar genes, as annotated by antiSMASH. In bold type, compounds empirically identified as produced by *Embleya australiensis* MST-111070 (by Microbial Screening Technologies Pty. Ltd., data not shown); in normal type, BGCs with more than 50% similar genes; in italics, BGCs with under 50% similar genes or no known similar BGC. Underlined, BGCs highly conserved and essential for the full biological cycle of streptomycetes. [5] Assigned based upon similarity to the known polyoxypeptin BGC (BGC0001036). [6] Identified by blastn search and PKS similarity: 92% identity with Figure 5 from patent US 7,288,396 B2. [7]Assigned by antiSMASH as most similar to SAL-2242 (BGC0000546) and SapB (BGC0000551) from *Streptomyces coelicolor* A3(2). [8] Highly conserved BGC among streptomycetes.

**Table S4: Highly conserved proteins between EEC1-like replicons.**

BlastP hits of proteins encoded by E. australiensis MST-11070 against *Embleya* sp. NBC 00888 and *Embleya* sp. NBC 00896 with an identity on both replicons >95 %. Conserved domains and predicted functions informed by NCBI's CDD search, and by work conducted in this manuscript. Clusters of conserved genes are highlighted in grey.

| Protein ID | %ID NBC00888 | %ID NBC00896 | Conserved Domains | Proposed Function |
|---|---|---|---|---|
| ctg2_731 | 95.4 | 96.3 | LarA | Cell envelope biosynthesis |
| ctg2_737 | 96 | 96 | PL-6 | Carbohydrate transport and metabolism |
| ctg2_738 | 95.6 | 97.2 | AfuC | Carbohydrate transport and metabolism |
| ctg2_739 | 97.6 | 97.2 | UgpE | Carbohydrate transport and metabolism |
| ctg2_740 | 97.8 | 98.1 | UgpA | Carbohydrate transport and metabolism |
| ctg2_827 | 97.2 | 97.2 | DppB | Transport |
| ctg2_1763 | 99.3 | 97.9 | Lrp | Regulation |
| ctg2_1888 | 100 | 99.3 | ParA | Replication |
| ctg2_1891 | 100 | 98.1 | Hypothetical | Replication |
| ctg2_2397 | 97.9 | 98.1 | HyuB | Biosynthesis |
| ctg2_2402 | 97.1 | 95.4 | FbaB | Carbohydrate transport and metabolism |
| ctg2_2511 | 96.6 | 96.6 | DppB | Transport |
| ctg2_2548 | 96 | 95.3 | IMPase | Phosphotase |
| ctg2_2987 | 95.4 | 95.5 | AraH | Carbohydrate transport and metabolism |
| ctg2_2990 | 95.8 | 96.6 | GlpR | Carbohydrate transport and metabolism |
| ctg2_3474 | 100 | 96.3 | large tegument protein UL36 | Unknown |

**Table S5: Features of *Embleya australiensis* MST-111070 EEC1 and large linear *Streptomyces* spp. replicons.**

Summary of biochemical and genetic composition of EEC1-3 compared to the largest *Streptomyces* megaplasmid reported to date pSCL4 from *Streptomyces clavuligerus*[59]. Spearman's and Pearson's correlations are shown.

| | EEC1 | EEC2 | EEC3 | pSCL4 |
|---|---|---|---|---|
| **Species** | *E. australiensis* | *E. australiensis* | *E. australiensis* | *S. clavuligerus* |
| **Strain** | MST-111070 | MST-111070 | MST-111070 | ATCC 27064 |
| **Classification** | Secondary Chromosome | Mega plasmid | Plasmid | Chromid |
| **Size** | 4.2 Mb | 0.3 MbX | 0.2 Mb | 1.8 Mb |
| **GC % (chromosomal)** | 71 % (71 %) | 69.1 (71 %) | 69.5 (71%) | 72 % (73%) |
| **Codon Bias (Correlation)** | Chromosomal (0.993/0.995) | Non-chromosomal (0.949/0.969) | Non-chromosomal (0.957/0.970) | Chromosomal (0.994/0.993) |
| **Functional Bias (Correlation)** | Chromosomal (0.972/0.984) | Non-chromosomal (0.543/0.456) | Non-chromosomal (0.499/0.325) | Chromosomal (0.938/0.917) |
| **BGC-density** | Higher than chromosome (~2x) | Higher than chromosome (~2x) | Higher than chromosome (~2x) | Higher than the chromosome (>2x) |
| **Replication System** | Unknown | Unknown | Plasmid | Plasmid |

### SUPPLEMENTARY FIGURES

**Figure S1. Identification and analysis of terminal inverted repeats.**

Blast analysis of the relationship between the terminal inverted repeats of the chromosomal contig (unitig_0) and the 4.2 Mb contig (unitig_1). A) Table of relevant blast HSPs; numbers in italics indicate that they are the last nucleotide of the contig, i.e. the HSP reaches the end of the contig. B) Graphical representation of the HSPs; each blast HSP is identified by a colour and number; the query and subject are represented by solid and dotted lines respectively.

**A**

| Hit No. | query id | subject id | % identity | alignment length | mismatches | gap opens | q. start | q. end | s. start | s. End |
|---|---|---|---|---|---|---|---|---|---|---|
| 1 | unitig_0 | unitig_0 | 99.99 | 21177 | 1 | 1 | 1 | 21176 | *7117762* | 7096586 |
| 2 | unitig_0 | unitig_1 | 99.94 | 33612 | 3 | 18 | 7084160 | *7117762* | 4184342 | *4217944* |
| 3 | unitig_0 | unitig_1 | 99.95 | 21184 | 1 | 9 | 1 | 21180 | 1 | 21179 |
| 4 | unitig_0 | unitig_1 | 99.95 | 21182 | 1 | 10 | 7096585 | *7117762* | 21176 | 1 |
| 5 | unitig_0 | unitig_1 | 99.95 | 21178 | 1 | 9 | 1 | 21176 | *4217944* | 4196774 |
| 6 | unitig_1 | unitig_1 | 99.91 | 21183 | 1 | 18 | 1 | 21176 | *4217944* | 4196773 |

**B**

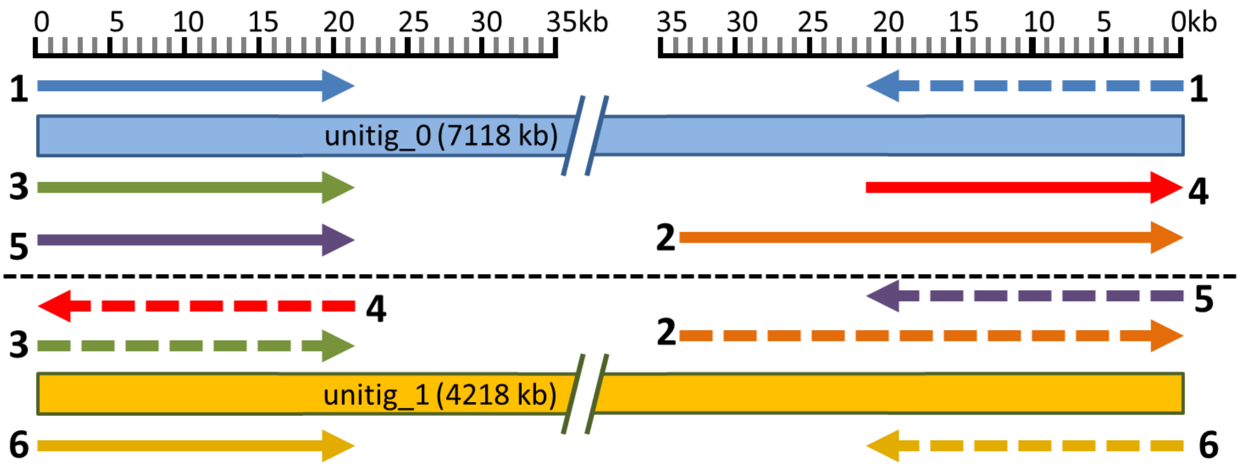

**Figure S2. Alignment of Bionano molecules and DNA sequence assembly.**

**A**. Overview of the alignment of BioNano molecules to the DNA sequence assembly, from left: chromosome, 4.2Mb contig (EEC1), the 302 kb linear EEC2, and circular plasmid EEC3. **B**. Full view of the alignment of BioNano molecules to the DNA sequence of EEC3, showing that because of the circular nature of this replicon, there are many Bionano molecules that extend outside the consensus DNA sequence used as reference, due to the different positions at which the plasmid molecule would have been opened during DNA extraction and BioNano assay.

**A**

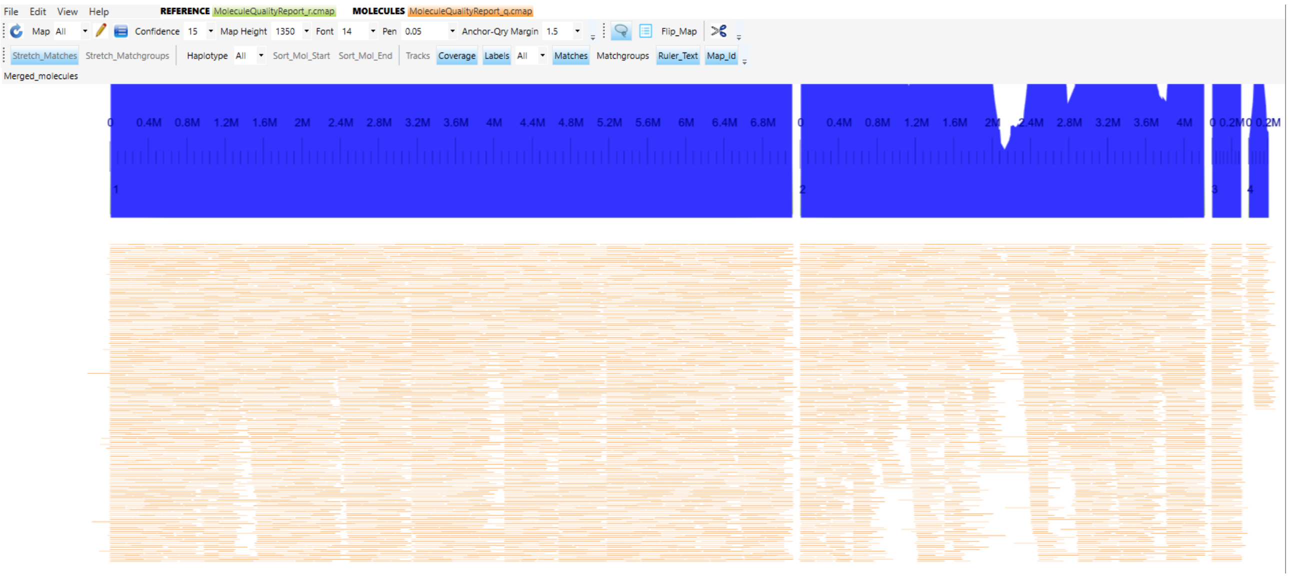

**B**

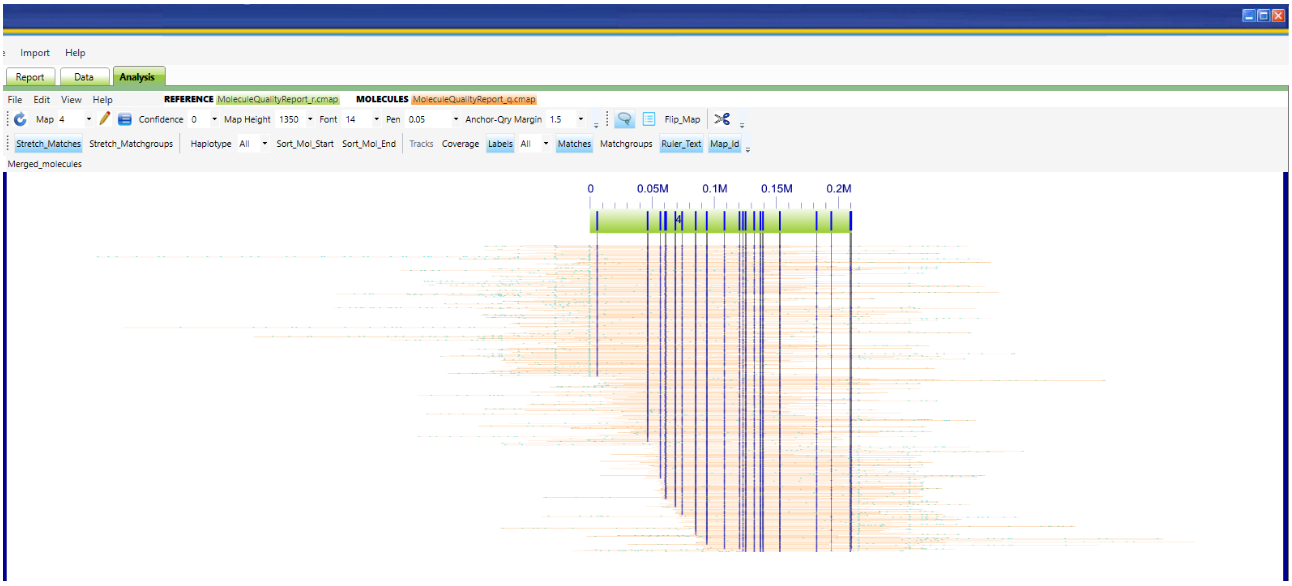

**Figure S3. Alignment of Bionano and sequence genome assemblies.**

Alignment of Bionano de-novo assembled contigs (blue) and DNA sequence assembly version 4 (green).

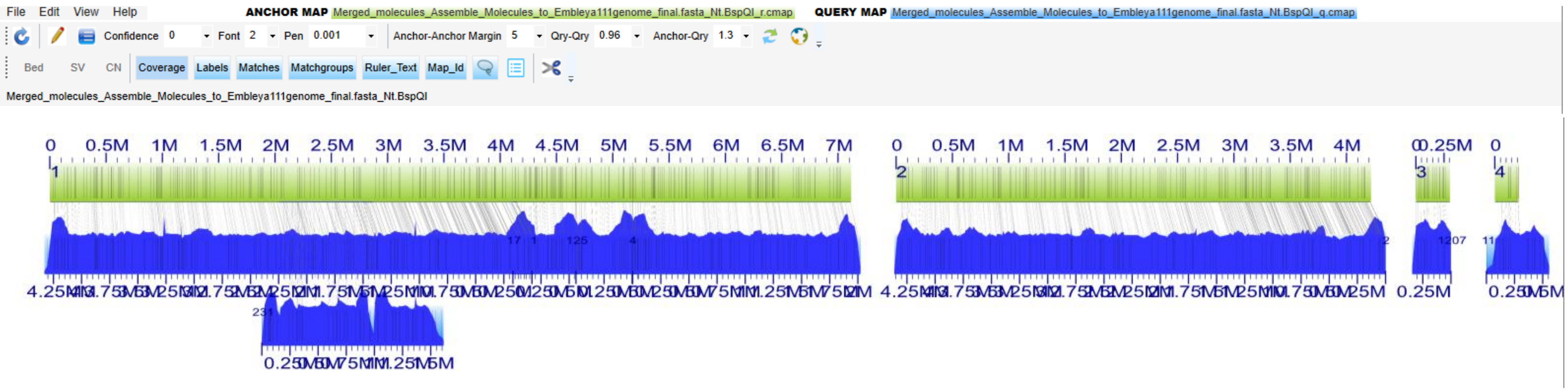

| | Match_Id | Anchor_Id | AnchorStart | AnchorEnd | Size | Qry_Id | Qry_Start | Qry_End | Orient | Confdn |
|---|---|---|---|---|---|---|---|---|---|---|
| 1 | 1 | 1 | 7,664 | 4,143,812 | 4,136,148 | 1 | 20 | 4,274,635 | - | 438.58 |
| 2 | 2 | 1 | 2,040,914 | 2,860,612 | 819,698 | 231 | 157,848 | 1,004,560 | + | 90.01 |
| 3 | 3 | 1 | 4,113,017 | 4,672,532 | 559,515 | 17 | 20 | 578,424 | + | 74.65 |
| 4 | 4 | 1 | 4,673,626 | 5,176,520 | 502,895 | 125 | 20 | 519,080 | + | 52.38 |
| 5 | 5 | 1 | 5,176,520 | 7,110,091 | 1,933,571 | 4 | 20 | 1,998,307 | + | 205.89 |
| 6 | 6 | 2 | 7,664 | 4,210,273 | 4,202,609 | 2 | 20 | 4,339,006 | - | 450.37 |
| 7 | 7 | 3 | 1,338 | 302,011 | 300,673 | 1207 | 20 | 310,441 | - | 36.4 |
| 8 | 8 | 4 | 5,647 | 210,490 | 204,843 | 11 | 80,320 | 291,203 | + | 21.86 |

Table of matches between Bionano de-novo assembled contigs and DNA-sequence assembly.

**Figure S4. Overview of the alignment of the 211 Kb EEC3 circular plasmid with its corresponding BioNano contig.**

The BioNano contig is more than double the size of the DNA-sequence contig; due to (and an indication of) the circular topology of this replicon, there are Bionano molecules that span in both directions of the purely artificial ends at which the circular sequence was opened (the stop codon of *parB*); it is also noticeable the repeated pattern of Bionano labels (vertical bars) in the Bionano contig (blue) at the right of the final match with the DNA-reference (green) as further support of the deducted circular topology.

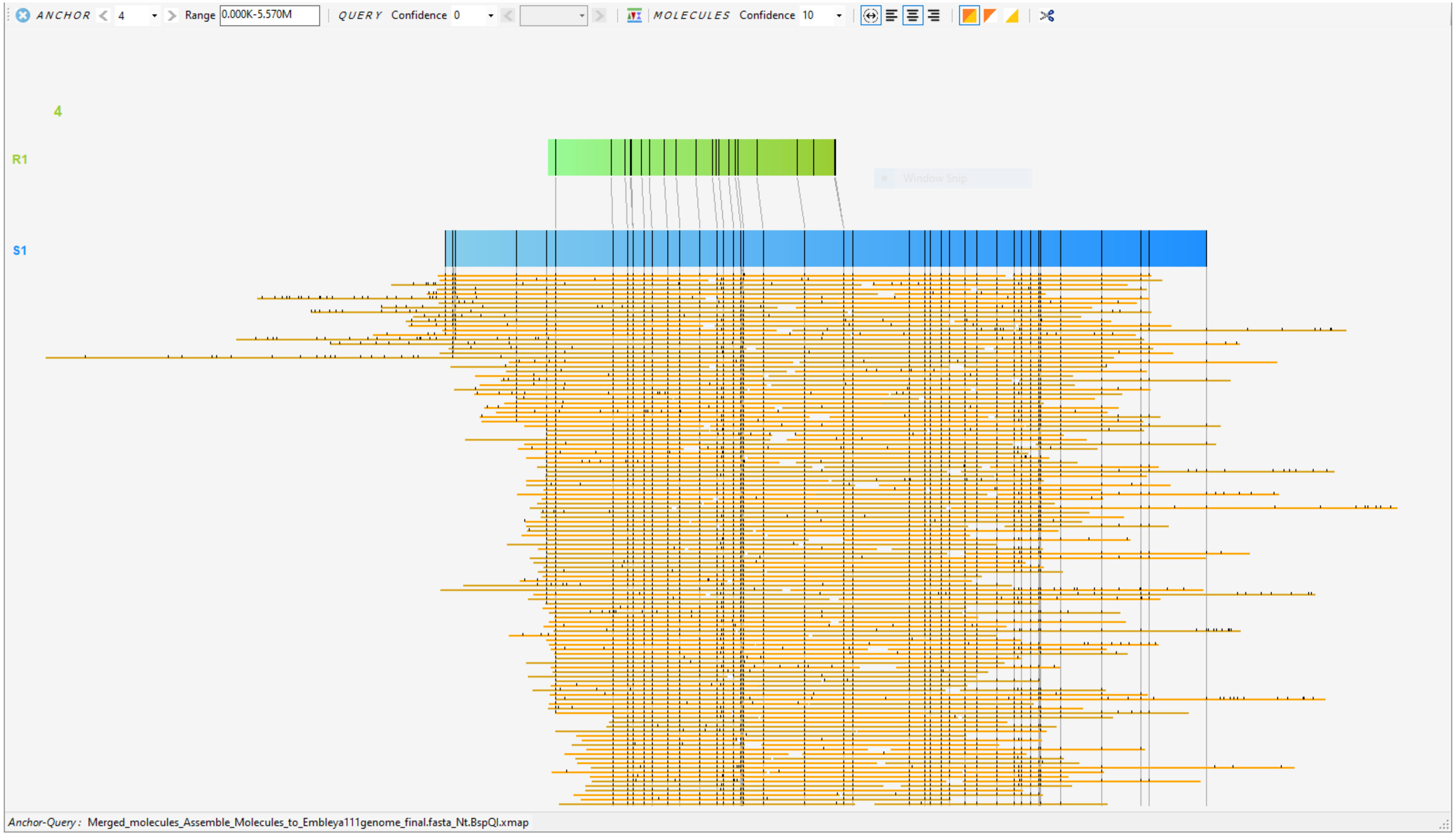

**Figure S5. Phylogenetic analysis provided by TYGS for full genome and 16S analysis of *Embleya australiensis* MST-111070.**

The final DNA assembly version 4 was uploaded to TYGS server [13] (on 2024-10-09 for the figures here shown). Phylogenetic tree based of full genome analysis (**A**) and 16S extended analysis offered by TYGS [13] with GGDC [15] (**B**) with only the genome of *E. australiensis* MST-111070 as query. This analysis initially revealed the membership of strain MST-111070 to the *Embleya* genus, and its close relationship with the genus *Yinghuangia*. (**A**) The values shown in blue are bootstrap values, in red are branch length. (**B**) The 16S tree was inferred under the GTR+CAT model and rooted by midpoint-rooting; the branches are scaled in terms of the expected number of substitutions per site; the numbers above the branches are support values when larger than 60% from Maximum likelihood (left) and maximum parsimony (right) bootstrapping. (modified from the automatic description provided by GGDC server). The *E. australiensis* MST-111070 genome is referred to as '111070v14genome_final' in the figure.

**A**

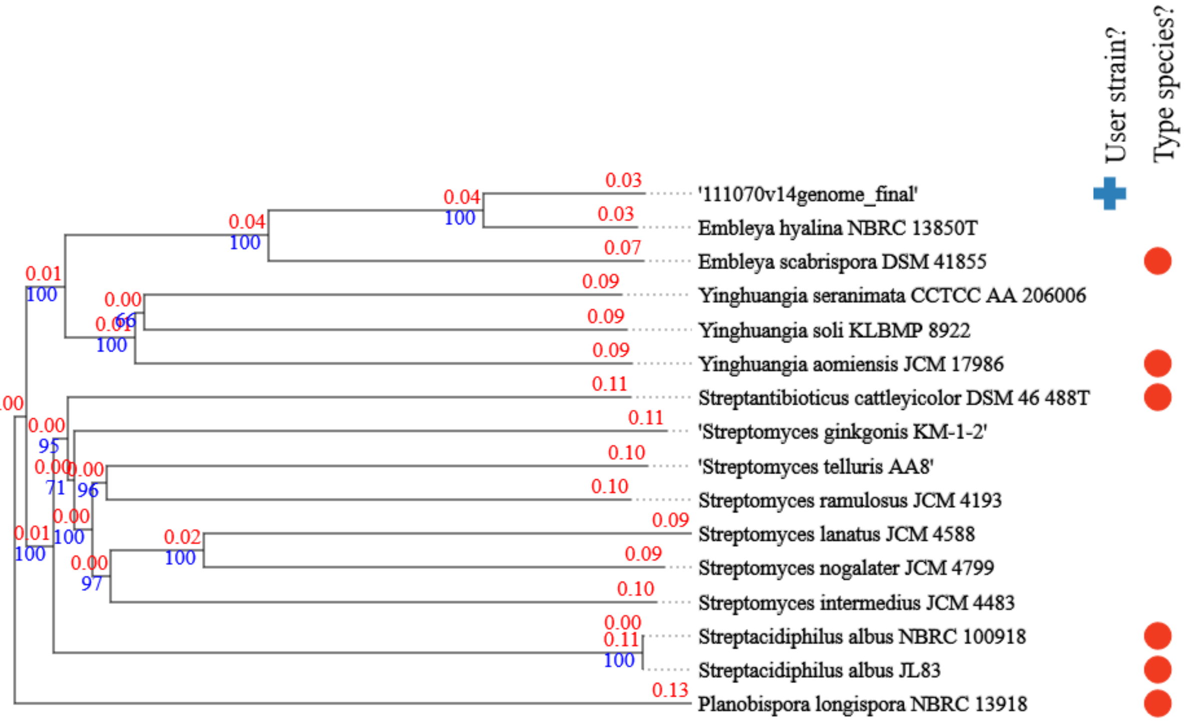

**B**

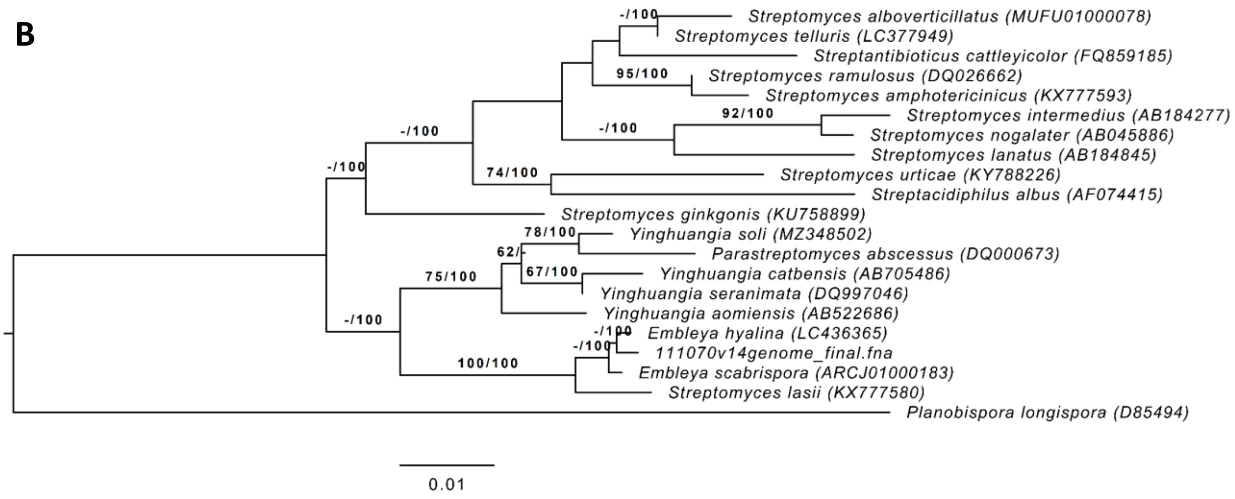

**Figure S6. Extended phylogenetic analysis provided by TYGS for full genome and 16S analysis of *Embleya australiensis* MST-111070.**

Next two pages.

The final DNA assembly version 4 (marked with red arrow) was uploaded to TYGS server [13] (on 2024-10-09 for the figures here shown) together with the genomes from NCBI databases identified as possibly belonging to *Embleya* and *Yinghuangia* genera (and other representative *Streptomyces* species). (**A**) Phylogenetic tree based of full genome analysis. Numbers above are bootstrap values. Blue cross at the right of the name indicates the genome as added as query by the user, while the red circle indicates that the strain is a type species of the genus. (**B**) 16S extended analysis offered by TYGS [13] with GGDC [15]. The tree was inferred under the GTR+CAT model and rooted by midpoint-rooting. The branches are scaled in terms of the expected number of substitutions per site. The numbers above the branches are support values when larger than 60% from Maximum likelihood (left) and maximum parsimony (right) bootstrapping. (modified from the automatic description provided by GGDC server).

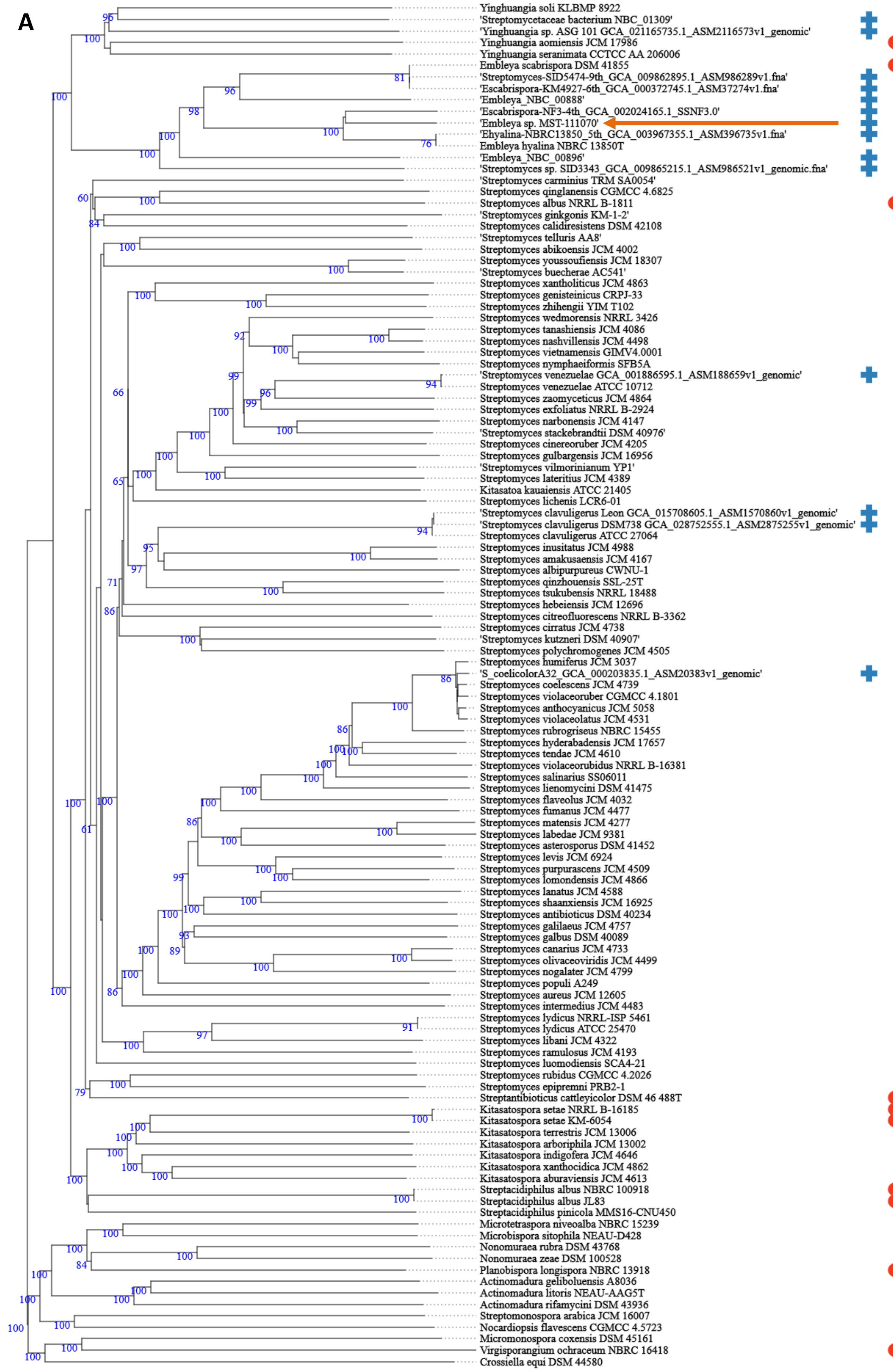
A
Yinghuangia soli KLBMP 8922
'Streptomycetaceae bacterium NBC_01309'
'Yinghuangia sp. ASG 101 GCA_021165735.1_ASM2116573v1_genomic'
Yinghuangia aomiensis JCM 17986
Yinghuangia seranimata CCTCC AA 206006
Embleya scabrispora DSM 41855
'Streptomyces-SID5474-9th_GCA_009862895.1_ASM986289v1.fna'
'Escabrispora-KM4927-6th_GCA_000372745.1_ASM37274v1.fna'
'Embleya_NBC_00888'
'Escabrispora-NF3-4th_GCA_002024165.1_SSNF3.0'
'Embleya sp. MST-111070'
'Ehyalina-NBRC13850_5th_GCA_003967355.1_ASM396735v1.fna'
Embleya hyalina NBRC 13850T
'Embleya_NBC_00896'
'Streptomyces sp. SID3343_GCA_009865215.1_ASM986521v1_genomic.fna'
'Streptomyces carminius TRM SA0054'
Streptomyces qinglanensis CGMCC 4.6825
Streptomyces albus NRRL B-1811
'Streptomyces ginkgonis KM-1-2'
Streptomyces calidiresistens DSM 42108
'Streptomyces telluris AA8'
Streptomyces abikoensis JCM 4002
Streptomyces youssoufiensis JCM 18307
'Streptomyces buecherae AC541'
Streptomyces xantholiticus JCM 4863
Streptomyces genisteinicus CRPJ-33
Streptomyces zhihengii YIM T102
Streptomyces wedmorensis NRRL 3426
Streptomyces tanashiensis JCM 4086
Streptomyces nashvillensis JCM 4498
Streptomyces vietnamensis GIMV4.0001
Streptomyces nymphaeiformis SFB5A
'Streptomyces venezuelae GCA_001886595.1_ASM188659v1_genomic'
Streptomyces venezuelae ATCC 10712
Streptomyces zaomyceticus JCM 4864
Streptomyces exfoliatus NRRL B-2924
Streptomyces narbonensis JCM 4147
'Streptomyces stackebrandtii DSM 40976'
Streptomyces cinereoruber JCM 4205
Streptomyces gulbargensis JCM 16956
'Streptomyces vilmorinianum YP1'
Streptomyces lateritius JCM 4389
Kitasatoa kauaiensis ATCC 21405
Streptomyces lichenis LCR6-01
'Streptomyces clavuligerus Leon GCA_015708605.1_ASM1570860v1_genomic'
'Streptomyces clavuligerus DSM738 GCA_028752555.1_ASM2875255v1_genomic'
Streptomyces clavuligerus ATCC 27064
Streptomyces inusitatus JCM 4988
Streptomyces amakusaensis JCM 4167
Streptomyces albipurpureus CWNU-1
Streptomyces qinzhouensis SSL-25T
Streptomyces tsukubensis NRRL 18488
Streptomyces hebeiensis JCM 12696
Streptomyces citreofluorescens NRRL B-3362
Streptomyces cirratus JCM 4738
'Streptomyces kutzneri DSM 40907'
Streptomyces polychromogenes JCM 4505
Streptomyces humiferus JCM 3037
'S_coelicolorA32_GCA_000203835.1_ASM20383v1_genomic'
Streptomyces coelescens JCM 4739
Streptomyces violaceoruber CGMCC 4.1801
Streptomyces anthocyanicus JCM 5058
Streptomyces violaceolatus JCM 4531
Streptomyces rubrogriseus NBRC 15455
Streptomyces hyderabadensis JCM 17657
Streptomyces tendae JCM 4610
Streptomyces violaceorubidus NRRL B-16381
Streptomyces salinarius SS06011
Streptomyces lienomycini DSM 41475
Streptomyces flaveolus JCM 4032
Streptomyces fumanus JCM 4477
Streptomyces matensis JCM 4277
Streptomyces labedae JCM 9381
Streptomyces asterosporus DSM 41452
Streptomyces levis JCM 6924
Streptomyces purpurascens JCM 4509
Streptomyces lomondensis JCM 4866
Streptomyces lanatus JCM 4588
Streptomyces shaanxiensis JCM 16925
Streptomyces antibioticus DSM 40234
Streptomyces galilaeus JCM 4757
Streptomyces galbus DSM 40089
Streptomyces canarius JCM 4733
Streptomyces olivaceoviridis JCM 4499
Streptomyces nogalater JCM 4799
Streptomyces populi A249
Streptomyces aureus JCM 12605
Streptomyces intermedius JCM 4483
Streptomyces lydicus NRRL-ISP 5461
Streptomyces lydicus ATCC 25470
Streptomyces libani JCM 4322
Streptomyces ramulosus JCM 4193
Streptomyces luomodiensis SCA4-21
Streptomyces rubidus CGMCC 4.2026
Streptomyces epipremni PRB2-1
Streptantibioticus cattleyicolor DSM 46 488T
Kitasatospora setae NRRL B-16185
Kitasatospora setae KM-6054
Kitasatospora terrestris JCM 13006
Kitasatospora arboriphila JCM 13002
Kitasatospora indigofera JCM 4646
Kitasatospora xanthocidica JCM 4862
Kitasatospora aburaviensis JCM 4613
Streptacidiphilus albus NBRC 100918
Streptacidiphilus albus JL83
Streptacidiphilus pinicola MMS16-CNU450
Microtetraspora niveoalba NBRC 15239
Microbispora sitophila NEAU-D428
Nonomuraea rubra DSM 43768
Nonomuraea zeae DSM 100528
Planobispora longispora NBRC 13918
Actinomadura geliboluensis A8036
Actinomadura litoris NEAU-AAG5T
Actinomadura rifamycini DSM 43936
Streptomonospora arabica JCM 16007
Nocardiopsis flavescens CGMCC 4.5723
Micromonospora coxensis DSM 45161
Virgisporangium ochraceum NBRC 16418
Crossiella equi DSM 44580

**B**

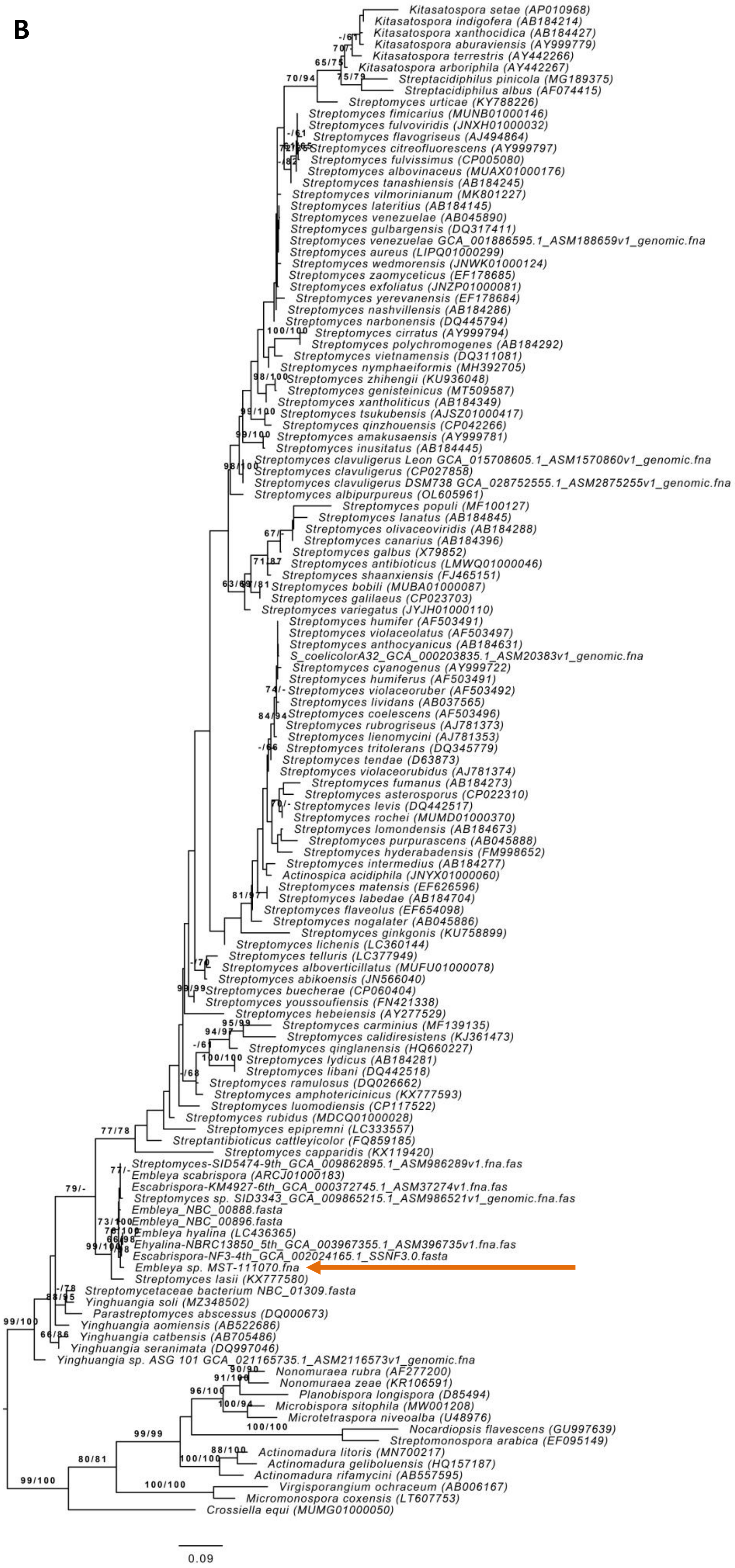
Kitasatospora setae (AP010968)
Kitasatospora indigofera (AB184214)
Kitasatospora xanthocidica (AB184427)
Kitasatospora aburaviensis (AY999779)
Kitasatospora terrestris (AY442266)
Kitasatospora arboriphila (AY442267)
Streptacidiphilus pinicola (MG189375)
Streptacidiphilus albus (AF074415)
Streptomyces urticae (KY788226)
Streptomyces fimicarius (MUNB01000146)
Streptomyces fulvoviridis (JNXH01000032)
Streptomyces flavogriseus (AJ494864)
Streptomyces citreofluorescens (AY999797)
Streptomyces fulvissimus (CP005080)
Streptomyces albovinaceus (MUAX01000176)
Streptomyces tanashiensis (AB184245)
Streptomyces vilmorinianum (MK801227)
Streptomyces lateritius (AB184145)
Streptomyces venezuelae (AB045890)
Streptomyces gulbargensis (DQ317411)
Streptomyces venezuelae GCA_001886595.1_ASM188659v1_genomic.fna
Streptomyces aureus (LIPQ01000299)
Streptomyces wedmorensis (JNWK01000124)
Streptomyces zaomyceticus (EF178685)
Streptomyces exfoliatus (JNZP01000081)
Streptomyces yerevanensis (EF178684)
Streptomyces nashvillensis (AB184286)
Streptomyces narbonensis (DQ445794)
Streptomyces cirratus (AY999794)
Streptomyces polychromogenes (AB184292)
Streptomyces vietnamensis (DQ311081)
Streptomyces nymphaeiformis (MH392705)
Streptomyces zhihengii (KU936048)
Streptomyces genisteinicus (MT509587)
Streptomyces xantholiticus (AB184349)
Streptomyces tsukubensis (AJSZ01000417)
Streptomyces qinzhouensis (CP042266)
Streptomyces amakusaensis (AY999781)
Streptomyces inusitatus (AB184445)
Streptomyces clavuligerus Leon GCA_015708605.1_ASM1570860v1_genomic.fna
Streptomyces clavuligerus (CP027858)
Streptomyces clavuligerus DSM738 GCA_028752555.1_ASM2875255v1_genomic.fna
Streptomyces albipurpureus (OL605961)
Streptomyces populi (MF100127)
Streptomyces lanatus (AB184845)
Streptomyces olivaceoviridis (AB184288)
Streptomyces canarius (AB184396)
Streptomyces galbus (X79852)
Streptomyces antibioticus (LMWQ01000046)
Streptomyces shaanxiensis (FJ465151)
Streptomyces bobili (MUBA01000087)
Streptomyces galilaeus (CP023703)
Streptomyces variegatus (JYJH01000110)
Streptomyces humifer (AF503491)
Streptomyces violaceolatus (AF503497)
Streptomyces anthocyanicus (AB184631)
S_coelicolorA32_GCA_000203835.1_ASM20383v1_genomic.fna
Streptomyces cyanogenus (AY999722)
Streptomyces humiferus (AF503491)
Streptomyces violaceoruber (AF503492)
Streptomyces lividans (AB037565)
Streptomyces coelescens (AF503496)
Streptomyces rubrogriseus (AJ781373)
Streptomyces lienomycini (AJ781353)
Streptomyces tritolerans (DQ345779)
Streptomyces tendae (D63873)
Streptomyces violaceorubidus (AJ781374)
Streptomyces fumanus (AB184273)
Streptomyces asterosporus (CP022310)
Streptomyces levis (DQ442517)
Streptomyces rochei (MUMD01000370)
Streptomyces lomondensis (AB184673)
Streptomyces purpurascens (AB045888)
Streptomyces hyderabadensis (FM998652)
Streptomyces intermedius (AB184277)
Actinospica acidiphila (JNYX01000060)
Streptomyces matensis (EF626596)
Streptomyces labedae (AB184704)
Streptomyces flaveolus (EF654098)
Streptomyces nogalater (AB045886)
Streptomyces ginkgonis (KU758899)
Streptomyces lichenis (LC360144)
Streptomyces telluris (LC377949)
Streptomyces alboverticillatus (MUFU01000078)
Streptomyces abikoensis (JN566040)
Streptomyces buecherae (CP060404)
Streptomyces youssoufiensis (FN421338)
Streptomyces hebeiensis (AY277529)
Streptomyces carminius (MF139135)
Streptomyces calidiresistens (KJ361473)
Streptomyces qinglanensis (HQ660227)
Streptomyces lydicus (AB184281)
Streptomyces libani (DQ442518)
Streptomyces ramulosus (DQ026662)
Streptomyces amphotericinicus (KX777593)
Streptomyces luomodiensis (CP117522)
Streptomyces rubidus (MDCQ01000028)
Streptomyces epipremni (LC333557)
Streptantibioticus cattleyicolor (FQ859185)
Streptomyces capparidis (KX119420)
Streptomyces-SID5474-9th_GCA_009862895.1_ASM986289v1.fna.fas
Embleya scabrispora (ARCJ01000183)
Escabrispora-KM4927-6th_GCA_000372745.1_ASM37274v1.fna.fas
Streptomyces sp. SID3343_GCA_009865215.1_ASM986521v1_genomic.fna.fas
Embleya_NBC_00888.fasta
Embleya_NBC_00896.fasta
Embleya hyalina (LC436365)
Ehyalina-NBRC13850_5th_GCA_003967355.1_ASM396735v1.fna.fas
Escabrispora-NF3-4th_GCA_002024165.1_SSNF3.0.fasta
Embleya sp. MST-111070.fna
Streptomyces lasii (KX777580)
Streptomycetaceae bacterium NBC_01309.fasta
Yinghuangia soli (MZ348502)
Parastreptomyces abscessus (DQ000673)
Yinghuangia aomiensis (AB522686)
Yinghuangia catbensis (AB705486)
Yinghuangia seranimata (DQ997046)
Yinghuangia sp. ASG 101 GCA_021165735.1_ASM2116573v1_genomic.fna
Nonomuraea rubra (AF277200)
Nonomuraea zeae (KR106591)
Planobispora longispora (D85494)
Microbispora sitophila (MW001208)
Microtetraspora niveoalba (U48976)
Nocardiopsis flavescens (GU997639)
Streptomonospora arabica (EF095149)
Actinomadura litoris (MN700217)
Actinomadura geliboluensis (HQ157187)
Actinomadura rifamycini (AB557595)
Virgisporangium ochraceum (AB006167)
Micromonospora coxensis (LT607753)
Crossiella equi (MUMG01000050)
70/94
65/75
75/79
100/100
98/100
99/100
77/78
79/-
73/100
99/100
99/99
80/81
100/100
0.09

**Figure S7. Heatmap and hierarchical clustering of codon usage correlation for secondary replicons in *S. clavuligerus* ATCC 27064.**

The figure was produced by Codoniser (this study). Correlation is calculated by Spearman's Rank.

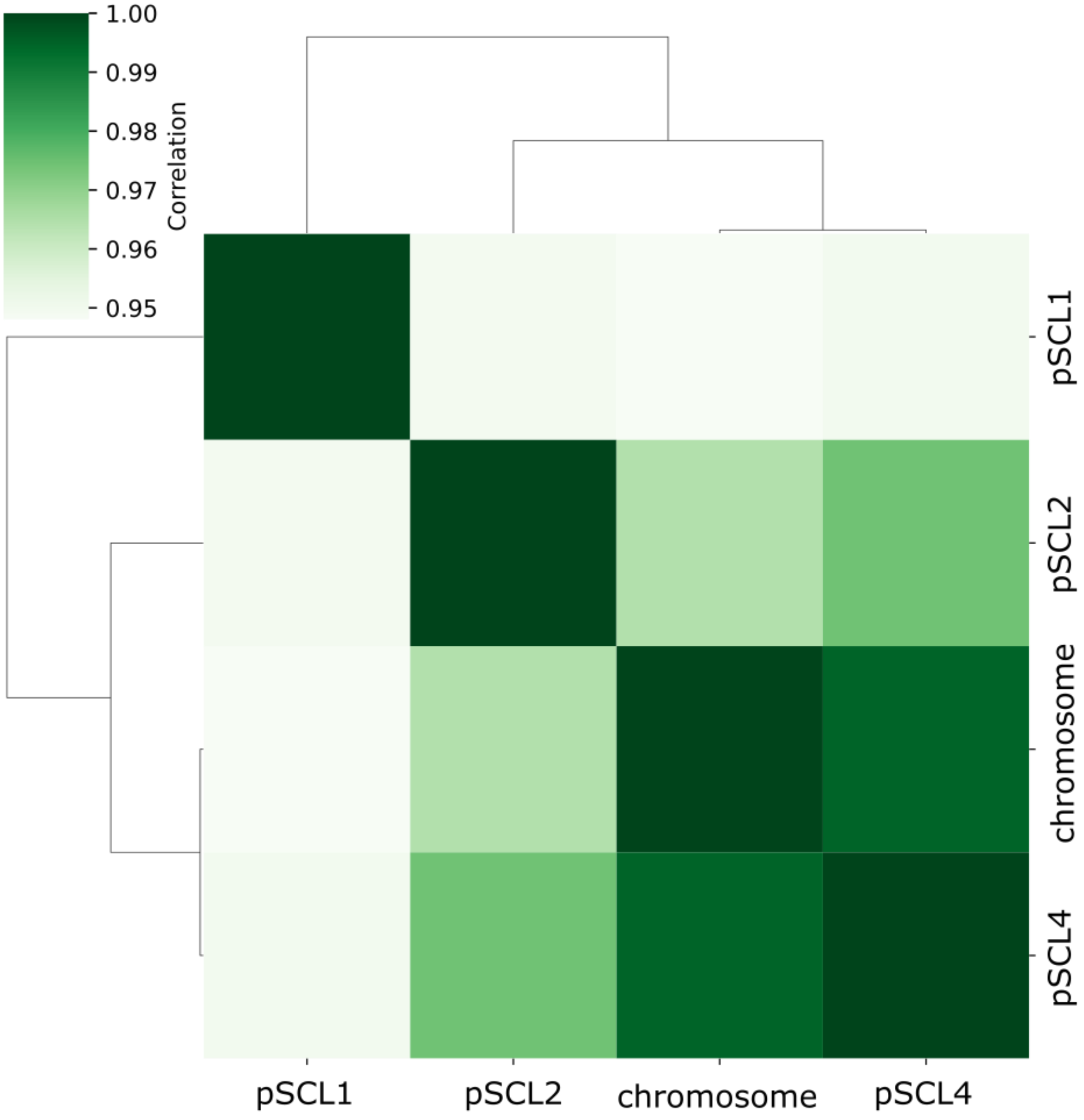

**Figure S8. Heatmap and hierarchical clustering of COG utilisation correlation for secondary replicons in *S. clavuligerus* ATCC 27064.**

The figure was produced by egger (this study). Correlation is calculated by Spearman's Rank.

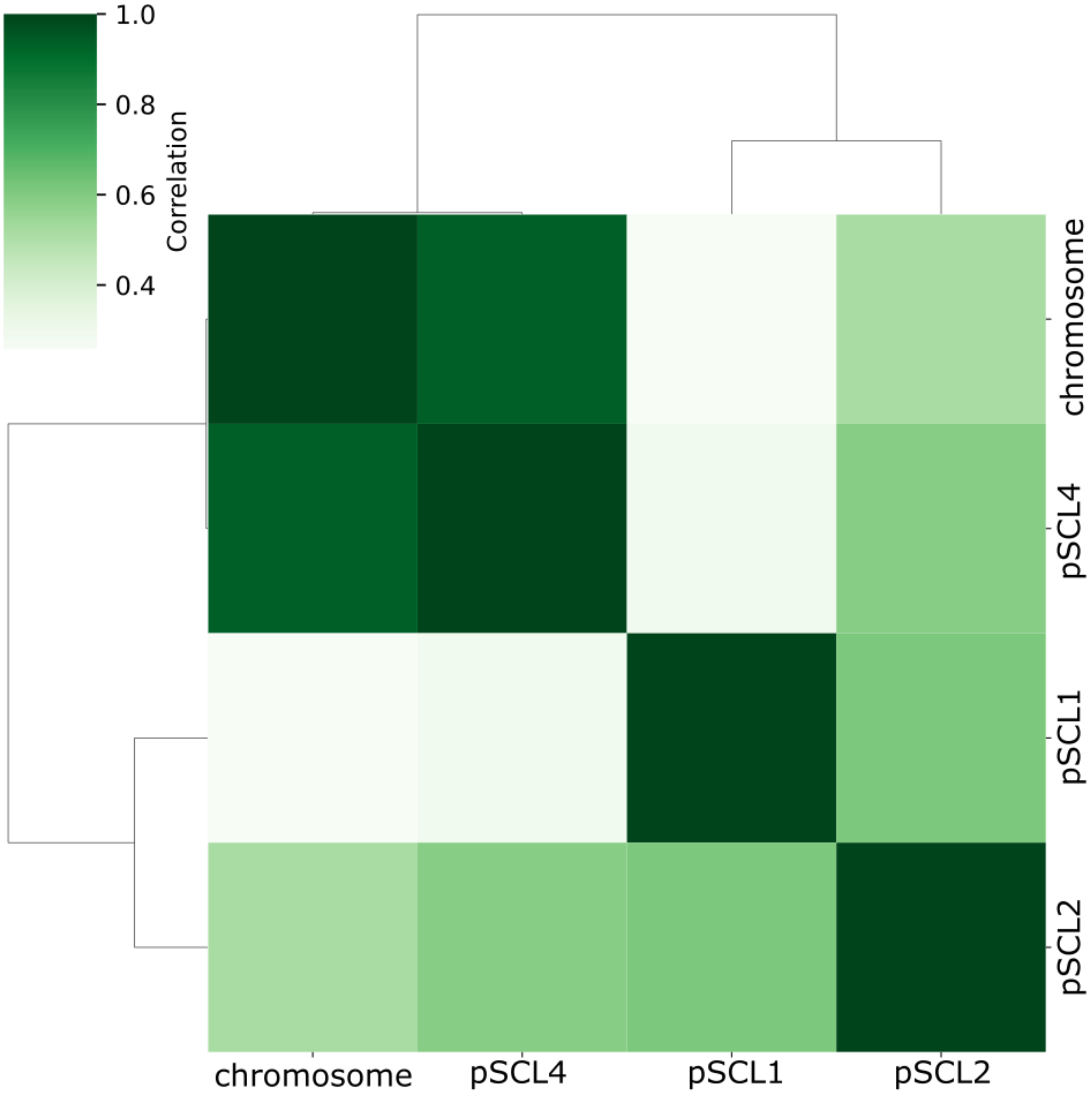

**Figure S9. Original output of the annotation of putative specialised metabolism biosynthetic gene clusters by antiSMASH**

Run on 2023-06-12 (continues on next two pages).

**unitig_0_Illu_ext_both_ends**

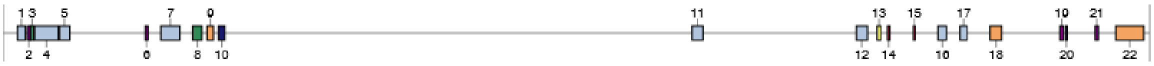

| Region | Type | From | To | Most similar known cluster | | Similarity |
|---|---|---|---|---|---|---|
| Region 1.1 | T1PKS, linaridin | 90,857 | 138,901 | legonaridin | RiPP | 41% |
| Region 1.2 | terpene | 150,651 | 171,443 | | | |
| Region 1.3 | CDPS | 171,621 | 192,325 | | | |
| Region 1.4 | NRPS-like, NRPS, T1PKS | 196,288 | 342,868 | incednine | Polyketide | 14% |
| Region 1.5 | thiopeptide, LAP, T1PKS | 349,551 | 414,110 | paromomycin | Saccharide | 7% |
| Region 1.6 | terpene | 882,668 | 901,353 | dudomycin A | NRP | 13% |
| Region 1.7 | indole, NRPS, T1PKS | 980,466 | 1,099,592 | polyoxypeptin | NRP+Polyketide | 75% |
| Region 1.8 | NRP-metallophore, NRPS | 1,177,888 | 1,235,978 | griseobactin | NRP | 92% |
| Region 1.9 | T3PKS | 1,266,421 | 1,307,485 | alkylresorcinol | Polyketide | 100% |
| Region 1.10 | other | 1,336,893 | 1,377,618 | himastatin | NRP | 12% |
| Region 1.11 | NRPS, ladderane | 4,277,131 | 4,344,826 | ishigamide | NRP+Polyketide | 100% |
| Region 1.12 | NAPAA, T1PKS, hglE-KS | 5,295,805 | 5,365,248 | hexacosalactone A | Other | 11% |
| Region 1.13 | lanthipeptide-class-iv | 5,425,093 | 5,448,404 | labyrinthopeptin A2/labyrinthopeptin A1/labyrinthopeptin A3 | RiPP:Lanthipeptide | 40% |
| Region 1.14 | NI-siderophore | 5,489,809 | 5,503,424 | schizokinen | Other | 30% |
| Region 1.15 | NI-siderophore | 5,649,743 | 5,661,680 | | | |
| Region 1.16 | terpene, betalactone | 5,803,706 | 5,856,397 | hopene | Terpene | 46% |
| Region 1.17 | NRPS-like, betalactone | 5,940,563 | 5,984,102 | nybomycin | Other | 55% |
| Region 1.18 | T1PKS | 6,127,601 | 6,199,142 | neoabyssomicin/abyssomicin | Polyketide | 31% |
| Region 1.19 | terpene | 6,559,664 | 6,581,427 | polyoxypeptin | NRP+Polyketide | 8% |
| Region 1.20 | RiPP-like | 6,595,549 | 6,606,349 | persiamycin A | Polyketide:Type II polyketide | 5% |
| Region 1.21 | terpene | 6,778,221 | 6,799,270 | | | |
| Region 1.22 | T1PKS | 6,905,975 | 7,081,035 | neoabyssomicin/abyssomicin | Polyketide | 50% |

unitig_1_Illu_Ext_both_ends

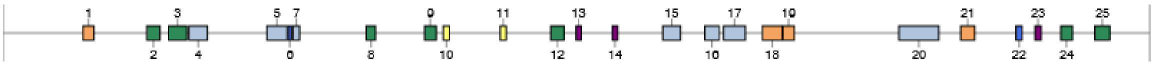

| Region | Type | From | To | Most similar known cluster | | Similarity |
|---|---|---|---|---|---|---|
| Region 2.1 | PKS-like | 294,513 | 335,535 | rustmicin | Polyketide:Iterative type I polyketide | 33% |
| Region 2.2 | NRPS, NRPS-like | 528,434 | 578,246 | WS9326 | NRP | 10% |
| Region 2.3 | NRPS, NRPS-like | 608,894 | 675,912 | himastatin | NRP | 56% |
| Region 2.4 | NRPS, betalactone | 683,679 | 752,046 | ohmyungsamycin A/ohmyungsamycin B | NRP | 12% |
| Region 2.5 | terpene, NRPS, aminocoumarin, ladderane, other, NRPS-like | 971,229 | 1,044,800 | acyldepsipeptide 1 | NRP+Polyketide | 26% |
| Region 2.6 | RiPP-like | 1,049,383 | 1,061,281 | | | |
| Region 2.7 | RRE-containing, butyrolactone | 1,066,340 | 1,091,725 | | | |
| Region 2.8 | NAPAA | 1,336,612 | 1,370,679 | γ-poly-L-2,4-diaminobutyric acid | NRP | 37% |
| Region 2.9 | NRPS | 1,550,635 | 1,593,139 | triacsin C | Other | 28% |
| Region 2.10 | lanthipeptide-class-iv | 1,619,382 | 1,641,973 | ulleungmycin | NRP | 5% |
| Region 2.11 | lanthipeptide-class-iii | 1,828,683 | 1,851,442 | SAL-2242 | RiPP:Lanthipeptide | 55% |
| Region 2.12 | NRPS | 2,015,110 | 2,063,485 | aurantimycin A | NRP+Polyketide | 5% |
| Region 2.13 | terpene | 2,107,635 | 2,127,304 | | | |
| Region 2.14 | terpene | 2,241,370 | 2,260,810 | | | |
| Region 2.15 | NRPS, T1PKS, NRPS-like | 2,426,374 | 2,492,278 | ECO-0501 | Polyketide | 19% |
| Region 2.16 | NRPS, lanthipeptide-class-ii | 2,581,031 | 2,634,701 | | | |
| Region 2.17 | NI-siderophore, NRPS, NRPS-like | 2,650,507 | 2,730,644 | peucechelin | NRP | 25% |
| Region 2.18 | T2PKS | 2,792,990 | 2,865,499 | spore pigment | Polyketide | 83% |
| Region 2.19 | T3PKS | 2,869,003 | 2,910,154 | R1128 | Polyketide | 21% |
| Region 2.20 | T1PKS, ladderane, arylpolyene, NRPS-like | 3,295,026 | 3,442,212 | akaeolide | Polyketide | 16% |
| Region 2.21 | hglE-KS | 3,522,775 | 3,573,744 | | | |
| Region 2.22 | lassopeptide | 3,725,618 | 3,748,424 | | | |
| Region 2.23 | terpene | 3,796,308 | 3,817,516 | ebelactone | Polyketide | 5% |
| Region 2.24 | NRPS-like | 3,890,646 | 3,933,120 | | | |
| Region 2.25 | NRPS-like | 4,015,181 | 4,071,452 | ecumicin | NRP | 10% |

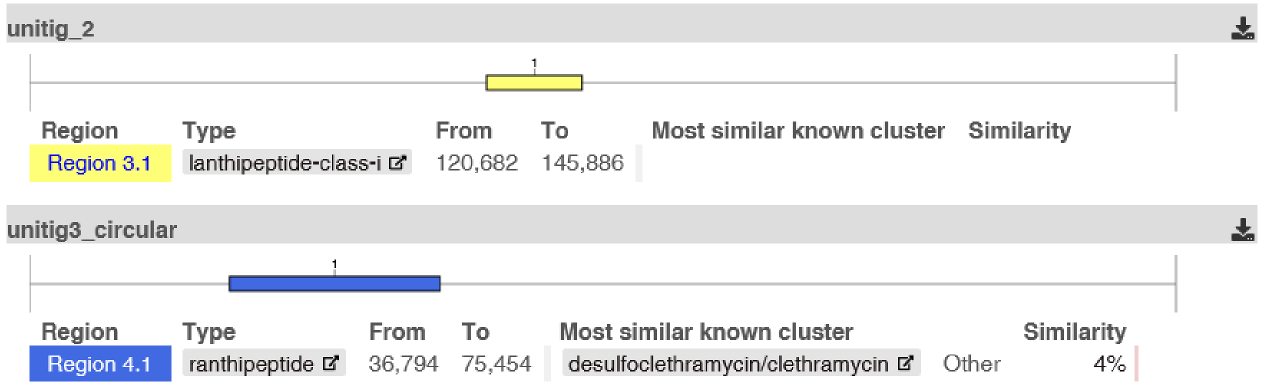

**Figure S10. Strain-specificity of proteins encoded by *E. australiensis* MST-11070 replicons at 70 % identity.**

Strip plots where each dot represents a single protein. Additional box plots are provided to show the median (red) and the quartile spread of the data points. All replicons encode a high number of strain-specific proteins, but it is accentuated in the plasmids EEC2 and EEC3.

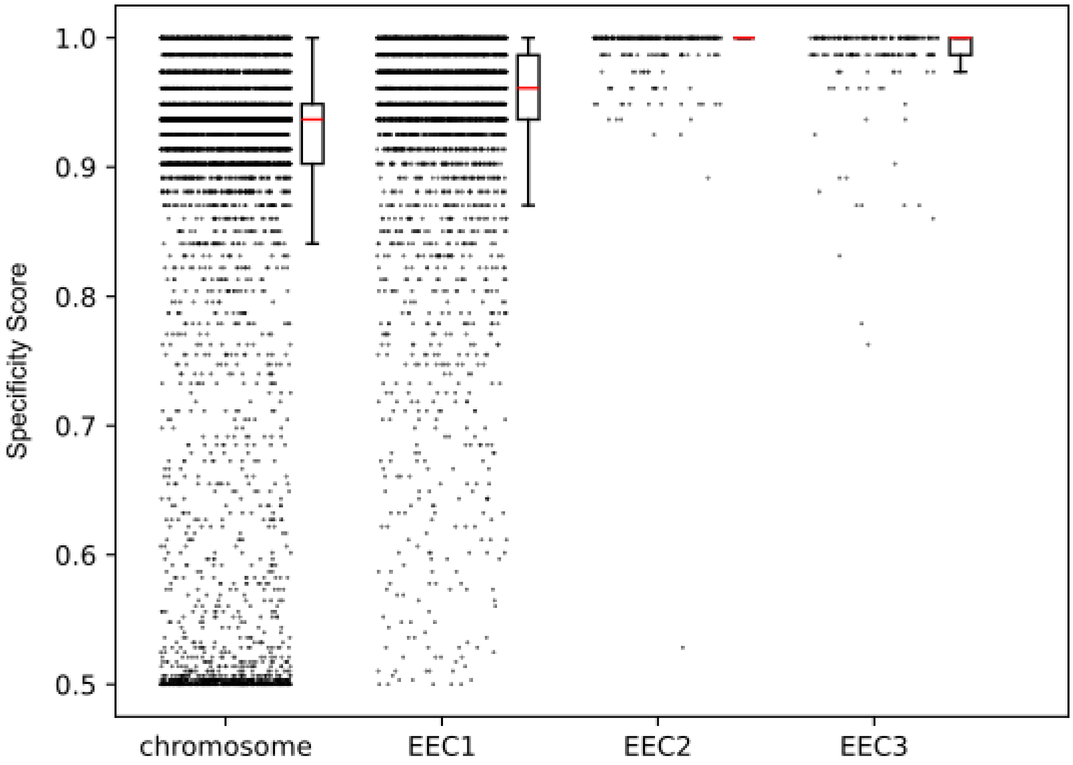

**Figure S11. Genus-specificity of proteins encoded by *E. australiensis* MST-11070 replicons at 70 % identity.**

Strip plots where each dot represents a single protein. Additional box plots are provided to show the median (red) and the quartile spread of the data points. Both EEC1 and the chromosome contain a large proportion of genus-specific genes, as opposed to the plasmids EEC2 and EEC3.

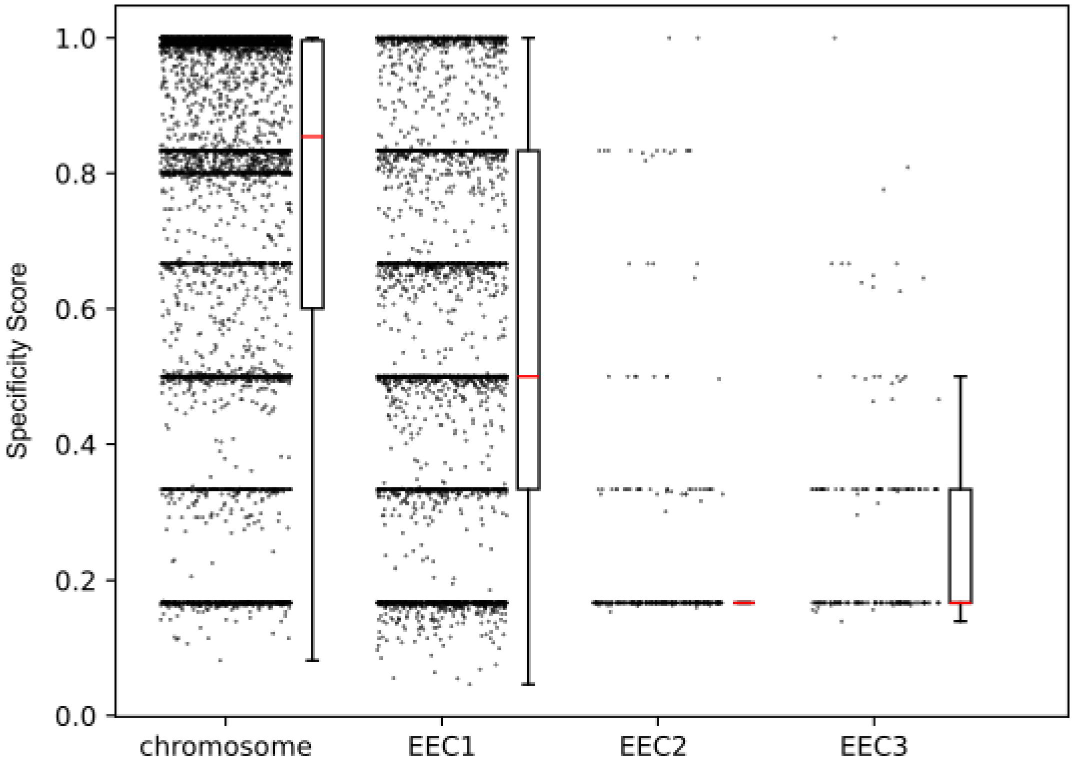

**Figure S12. Family-specificity of proteins encoded by *E. australiensis* MST-11070 replicons at 70 % identity.**

Strip plots where each dot represents a single protein. Additional box plots are provided to show the median (red) and the quartile spread of the data points. Specificity scores drop considerably at the family level, although notably, the chromosome still contains some highly family-specific proteins.

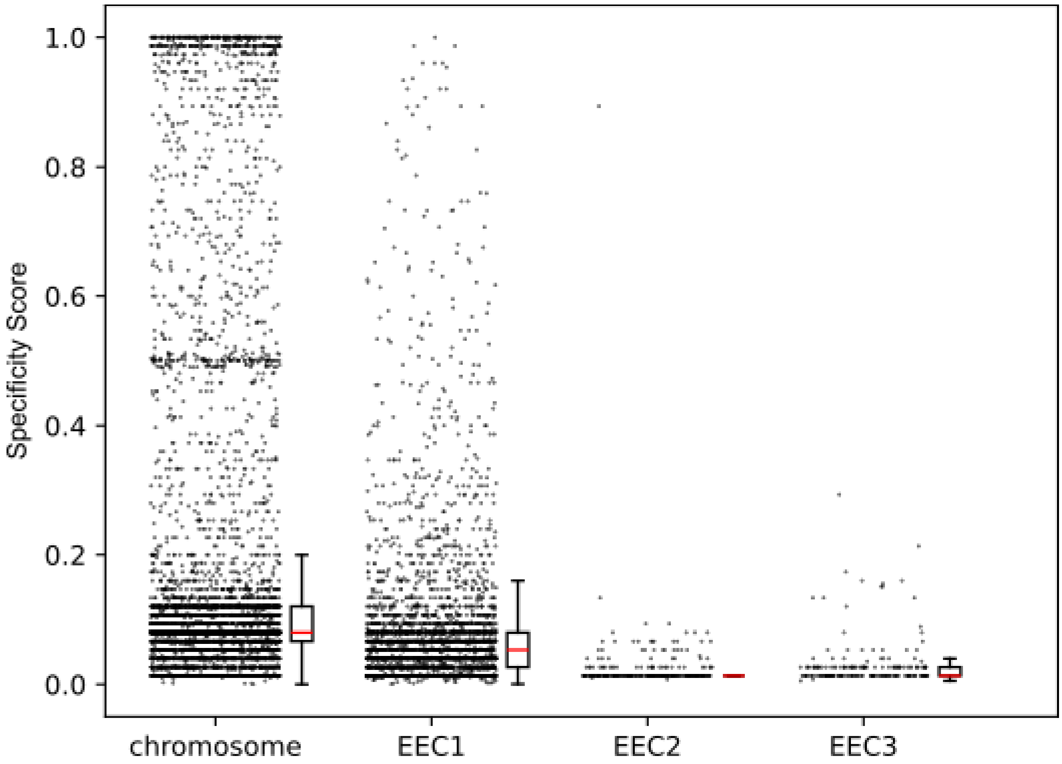

**Figure S13. Alignment of *Embleya australiensis* MST-111070 EEC1 with large secondary replicons from other *Embleya* species.**

Dot-plots showing the similarity and synteny of the secondary replicon EEC1 from *E. australiensis* MST-111070 with large secondary replicons identified in high quality genome assemblies from two additional *Embleya species*.

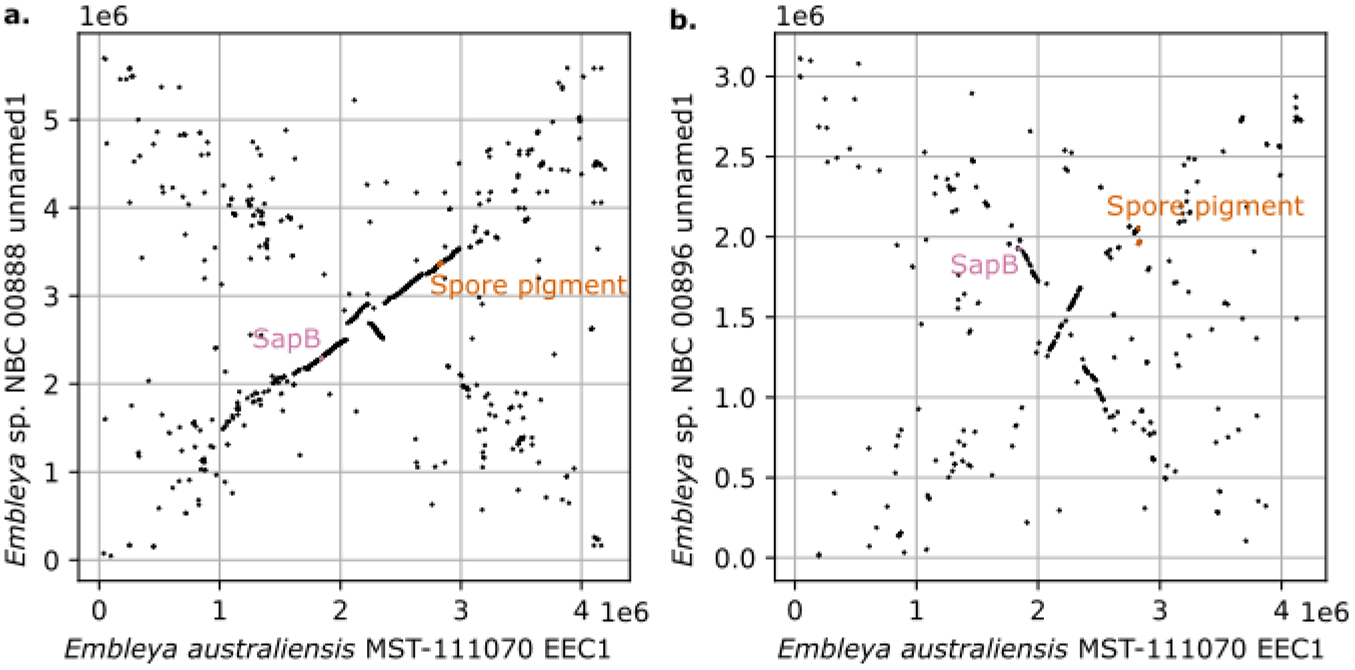

**Figure S14. antiSMASH annotation of the large secondary replicon of high-quality genome assemblies of other *Embleya* species.**

View of the antiSMASH annotation of the large secondary replicon of *Embleya* sp. NBC 00888 (5.7 Mb, top) and *Embleya* sp. NBC 00896 (3.1 Mb, bottom), showing the presence of the highly conserved and essential BGCs for the spore pigment and SapB (not readily identified by antiSMASH in NCB 00896 in any of its replicons) in this secondary replicon, instead of in the chromosome as it is the case in all other genera of the *Streptomycetaceae* family. For the analysis, we uploaded to antiSMASH the annotated NCBI RefSeq assembly files GCF_045207655.1 and GCF_045207785.1.

**NZ_CP108785.1** (Embleya sp. NBC_00888)

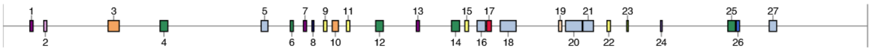

| Region | Type | From | To | Most similar known cluster | | Similarity |
|---|---|---|---|---|---|---|
| Region 2.1 | terpene | 179,678 | 200,661 | mycotrienin I | NRP+Polyketide | 7% |
| Region 2.2 | phenazine | 271,236 | 291,739 | pyocyanine | Other | 71% |
| Region 2.3 | T1PKS | 697,343 | 771,720 | spectinabilin/orinocin/SNF4435C/ SNF4435D | Polyketide:Modular type I polyketide | 90% |
| Region 2.4 | NRPS-like , NRPS | 1,042,149 | 1,093,548 | | | |
| Region 2.5 | arylpolyene , NRPS-like | 1,712,211 | 1,757,005 | calicheamicin | Polyketide | 8% |
| Region 2.6 | CDPS | 1,904,899 | 1,925,612 | | | |
| Region 2.7 | terpene | 1,992,235 | 2,013,305 | ishigamide | NRP+Polyketide | 11% |
| Region 2.8 | hydrogen-cyanide | 2,049,038 | 2,062,163 | | | |
| Region 2.9 | lanthipeptide-class-iv | 2,125,344 | 2,147,956 | ulleungmycin | NRP | 5% |
| Region 2.10 | T3PKS | 2,183,139 | 2,224,251 | napyradiomycin | Terpene | 7% |
| Region 2.11 | lanthipeptide-class-iii | 2,277,354 | 2,299,966 | SapB | RiPP:Lanthipeptide | 75% |
| Region 2.12 | NRPS | 2,471,282 | 2,521,702 | | | |
| Region 2.13 | terpene | 2,740,358 | 2,761,458 | | | |
| Region 2.14 | NRPS | 2,973,699 | 3,027,354 | stenothricin | NRP:Cyclic depsipeptide | 9% |
| Region 2.15 | lanthipeptide-class-i | 3,062,251 | 3,086,787 | | | |
| Region 2.16 | NRPS , lanthipeptide-class-ii | 3,143,339 | 3,198,749 | leinamycin | NRP+Polyketide:Modular type I polyketide+Polyketide:Trans-AT type I polyketide | 4% |
| Region 2.17 | NI-siderophore | 3,206,430 | 3,238,052 | peucechelin | NRP | 20% |
| Region 2.18 | NRPS-like , T2PKS | 3,296,406 | 3,401,918 | spore pigment | Polyketide | 83% |
| Region 2.19 | indole | 3,683,838 | 3,704,992 | | | |
| Region 2.20 | NRP-metallophore , NRPS , terpene , NAPAA | 3,729,075 | 3,839,439 | madurastatin A2/madurastatin E1/ madurastatin F/madurastatin G1/ madurastatin A1 | NRP | 47% |
| Region 2.21 | hglE-KS , NRPS | 3,844,397 | 3,914,475 | triacsin C | Other | 28% |
| Region 2.22 | lanthipeptide-class-iv | 4,004,293 | 4,026,980 | | | |
| Region 2.23 | ectoine | 4,135,088 | 4,145,486 | ectoine | Other | 100% |
| Region 2.24 | butyrolactone | 4,359,055 | 4,369,978 | | | |
| Region 2.25 | NRPS | 4,804,906 | 4,855,300 | bonnevillamide D/bonnevillamide E | NRP | 10% |
| Region 2.26 | linaridin | 4,861,744 | 4,882,661 | legonaridin | RiPP | 41% |
| Region 2.27 | terpene , NRPS , T1PKS | 5,079,299 | 5,128,694 | dutomycin | Polyketide | 4% |

**NZ_CP108777.1** (Embleya sp. NBC_00896)

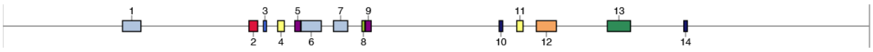

| Region | Type | From | To | Most similar known cluster | | Similarity |
|---|---|---|---|---|---|---|
| Region 2.1 | T3PKS , NRPS | 433,330 | 499,867 | feglymycin | NRP | 36% |
| Region 2.2 | NI-siderophore | 890,414 | 922,039 | peucechelin | NRP | 20% |
| Region 2.3 | RiPP-like | 942,644 | 953,840 | | | |
| Region 2.4 | lanthipeptide-class-i | 993,965 | 1,018,408 | | | |
| Region 2.5 | terpene | 1,056,104 | 1,077,156 | | | |
| Region 2.6 | terpene , NRPS | 1,078,830 | 1,151,878 | neocarazostatin A | Other | 60% |
| Region 2.7 | T3PKS , NAPAA | 1,195,911 | 1,247,420 | napyradiomycin | Terpene | 7% |
| Region 2.8 | ectoine | 1,298,872 | 1,309,270 | ectoine | Other | 100% |
| Region 2.9 | terpene | 1,311,164 | 1,332,270 | | | |
| Region 2.10 | hydrogen-cyanide | 1,794,792 | 1,807,731 | aborycin | RiPP | 21% |
| Region 2.11 | lanthipeptide-class-ii | 1,858,317 | 1,881,226 | | | |
| Region 2.12 | T2PKS | 1,929,028 | 2,001,585 | spore pigment | Polyketide | 83% |
| Region 2.13 | NRPS , NRP-metallophore | 2,185,839 | 2,269,601 | madurastatin D1/madurastatin D2/(-)-Madurastatin C1 | NRP | 8% |
| Region 2.14 | hydrogen-cyanide | 2,462,196 | 2,475,223 | | | |

**Figure S15. Artemis Comparison Tool view of the genome sequence comparison.**

Artemis Comparison Tool view of the genome sequence comparison between (**A**) *Embleya australiensis* MST-111070 and *Embleya scabrispora* NF3 (minimum Score cut-off of 3795 was set to facilitate visualisation of most relevant similarities) and (**B**) strain 111070 and *Embleya hyalina* (minimum Score cut-off of 3344).

**A** **B**

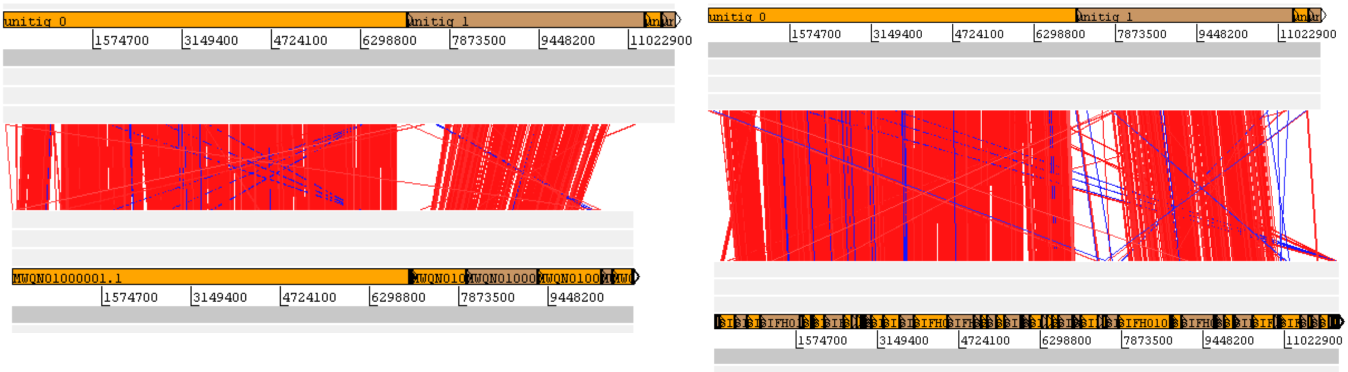

**Figure S16. Alignment of the genome assemblies of *Embleya australiensis* MST-111070 with available genome assemblies from other *Embleya* strains.**

Alignment with Mauve of the genome assemblies of *E. australiensis* MST-111070 with all the available genome assemblies for *Embleya* strains. The "backbone colour" representation has been selected, meaning that similar genome regions shared across all genomes are highlighted with mauve colour, while regions shared between two or three are marked with other colours. Most of the *E. australiensis* MST-111070 genome is shared across all three other assemblies, that includes most of the 4.2 Mb contig, indicating that this contig is not just a strain or species-specific extrachromosomal replicon but a genus shared characteristic.

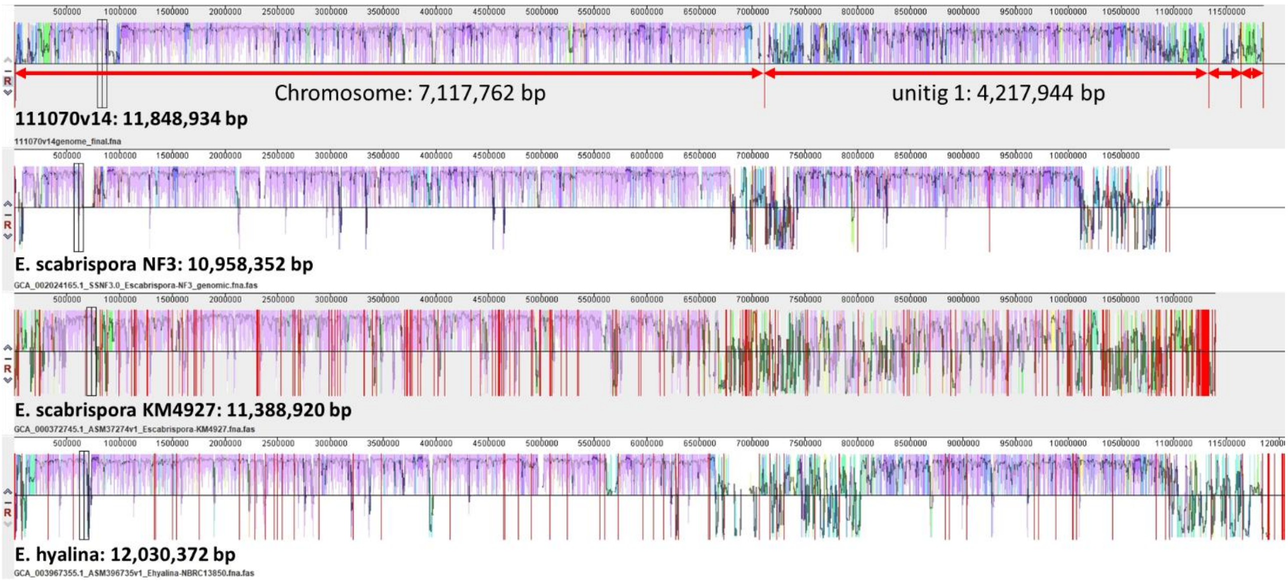

**REFERENCES**

1. **Rutherford K, Parkhill J, Crook J, Horsnell T, Rice P, Rajandream M-A, Barrell B**. Artemis: sequence visualization and annotation. *Bioinformatics* 2000;16:944–945, doi:10.1093/bioinformatics/16.10.944.

2. **Carver TJ, Rutherford KM, Berriman M, Rajandream M-A, Barrell BG, Parkhill J**. ACT: the Artemis comparison tool. *Bioinformatics* 2005;21:3422–3423, doi:10.1093/bioinformatics/bti553.

3. **Staden R**. The staden sequence analysis package. *Mol Biotechnol* 1996;5:233, doi:10.1007/BF02900361.

4. **Staden R, Beal KF, Bonfield JK**. The Staden Package, 1998. In: Misener S, Krawetz SA (editors). *Bioinformatics methods and protocols*. Totowa, NJ: Humana Press. pp. 115–130 ISBN 978-1-59259-192-3.

5. **Altschul SF, Madden TL, Schäffer AA, Zhang J, Zhang Z, Miller W, Lipman DJ**. Gapped BLAST and PSI-BLAST: a new generation of protein database search programs. *Nucleic Acids Res* 1997;25:3389–3402, doi:10.1093/nar/25.17.3389.

6. **Santiago-Sotelo P, Ramirez-Prado JH**. prfectBLAST: a platform-independent portable front end for the command terminal BLAST+ stand-alone suite. *BioTechniques* 2012;53:299–300, doi:10.2144/000113953.

7. **Blin K, Shaw S, Augustijn HE, Reitz ZL, Biermann F, Alanjary M, Fetter A, Terlouw BR, Metcalf WW, Helfrich EJN, van Wezel GP, Medema MH, Weber T**. antiSMASH 7.0: new and improved predictions for detection, regulation, chemical structures and visualisation. *Nucleic Acids Research* 2023;gkad344, doi:10.1093/nar/gkad344.

8. **Aziz RK, Bartels D, Best AA, DeJongh M, Disz T, Edwards RA, Formsma K, Gerdes S, Glass EM, Kubal M, Meyer F, Olsen GJ, Olson R, Osterman AL, Overbeek RA, McNeil LK, Paarmann D, Paczian T, Parrello B, Pusch GD, Reich C, Stevens R, Vassieva O, Vonstein V, Wilke A, Zagnitko O**. The RAST Server: Rapid Annotations using Subsystems Technology. *BMC Genomics* 2008;9:75, doi:10.1186/1471-2164-9-75.

9. **Overbeek R, Olson R, Pusch GD, Olsen GJ, Davis JJ, Disz T, Edwards RA, Gerdes S, Parrello B, Shukla M, Vonstein V, Wattam AR, Xia F, Stevens R**. The SEED and the Rapid Annotation of microbial genomes using Subsystems Technology (RAST). *Nucl Acids Res* 2014;42:D206–D214, doi:10.1093/nar/gkt1226.

10. **Hyatt D, Chen G-L, Locascio PF, Land ML, Larimer FW, Hauser LJ**. Prodigal: prokaryotic gene recognition and translation initiation site identification. *BMC Bioinformatics* 2010;11:119, doi:10.1186/1471-2105-11-119.

11. **Darling AE, Mau B, Perna NT**. progressiveMauve: Multiple genome alignment with gene gain, loss and rearrangement. *PLOS ONE* 2010;5:e11147, doi:10.1371/journal.pone.0011147.

12. **Simão FA, Waterhouse RM, Ioannidis P, Kriventseva EV, Zdobnov EM**. BUSCO: assessing genome assembly and annotation completeness with single-copy orthologs. *Bioinformatics* 2015;31:3210–3212, doi:10.1093/bioinformatics/btv351.

13. **Meier-Kolthoff JP, Göker M**. TYGS is an automated high-throughput platform for state-of-the-art genome-based taxonomy. *Nature Communications* 2019;10:2182, doi:10.1038/s41467-019-10210-3.

14. **Meier-Kolthoff JP, Auch AF, Klenk H-P, Göker M**. Genome sequence-based species delimitation with confidence intervals and improved distance functions. *BMC Bioinformatics* 2013;14:60, doi:10.1186/1471-2105-14-60.

15. **Meier-Kolthoff JP, Carbasse JS, Peinado-Olarte RL, Göker M**. TYGS and LPSN: a database tandem for fast and reliable genome-based classification and nomenclature of prokaryotes. *Nucleic Acids Research* 2022;50:D801–D807, doi:10.1093/nar/gkab902.

16. **Buchfink B, Xie C, Huson DH**. Fast and sensitive protein alignment using DIAMOND. *Nat Methods* 2015;12:59–60, doi:10.1038/nmeth.3176.
